# Implications of taxonomic bias for microbial differential-abundance analysis

**DOI:** 10.1101/2022.08.19.504330

**Authors:** Michael R. McLaren, Jacob T. Nearing, Amy D. Willis, Karen G. Lloyd, Benjamin J. Callahan

## Abstract

Differential-abundance (DA) analyses enable microbiome researchers to assess how microbial species vary in relative or absolute abundance with specific host or environmental conditions, such as health status or pH. These analyses typically use sequencing-based community measurements that are taxonomically biased to measure some species more efficiently than others. Understanding the effects that taxonomic bias has on the results of a DA analysis is essential for achieving reliable and translatable findings; yet currently, these effects are unknown. Here, we characterized these effects for DA analyses of both relative and absolute abundances, using a combination of mathematical theory and data analysis of real and simulated case studies. We found that, for analyses based on species proportions, taxonomic bias can cause significant errors in DA results if the average measurement efficiency of the community is associated with the condition of interest. These errors can be avoided by using more robust DA methods (based on species ratios) or quantified and corrected using appropriate controls. Wide adoption of our recommendations can improve the reproducibility, interpretability, and translatability of microbiome DA studies.

This manuscript was rendered from commit 7412a36 of https://github.com/mikemc/differential-abundance-theory. Supporting data analyses can be found in the accompanying computational research notebook. Please post comments or questions on GitHub. The manuscript is licensed under a CC BY 4.0 License. See the GitHub Releases or Zenodo record for earlier versions.

## 1 Introduction

One of the most basic questions we can ask about microbial communities is: How do different microbial taxa vary in abundance—across space, time, and host or environmental conditions? Marker-gene and shotgun metagenomic sequencing (jointly, MGS) can be used to measure the abundances of thousands of species simultaneously, making it possible to ask this question on a community-wide scale. In these *differential-abundance (DA) analyses*, the change in abundance of a microbial taxon across samples or conditions is used to infer ecological dynamics or find microbes that are associated with specific host diseases or environmental conditions. Standard MGS measurements lose information about total microbial density and so are typically used to analyze the abundances of taxa relative to each other (*relative abundances*). But new methods are increasingly used to provide absolute information, making it possible to analyze changes in absolute cell density, biomass, or genome copy number (*absolute abundances*). In its various forms, DA is among the most common analyses applied to MGS data.

Unfortunately, these DA analyses are built on a fundamentally flawed foundation.

MGS measurements are *taxonomically biased*: Microbial species vary dramatically in how efficiently they are measured—that is, converted from cells into taxonomically classified sequencing reads—by a given MGS protocol (McLaren, Willis, and Callahan (2019)). This bias arises from variation in how species respond to each step in an MGS protocol, from sample collection to bioinformatic classification. Although often associated with features specific to marker-gene sequencing—the variation among species in marker copy numbers and in primer-binding and amplification efficiencies—the existence of large variation in DNA extraction efficiencies and in the ability to correctly classify reads make taxonomic bias a universal feature of both marker-gene and shotgun measurements. As a result, MGS measurements provide inaccurate representations of actual community composition and tend to differ systematically across protocols, studies, and even experimental batches within a study (Yeh et al. (2018), McLaren, Willis, and Callahan (2019)). These errors can supersede sizable biological differences (e.g. Lozupone et al. (2013)) and may have contributed to failed replications of prominent DA results such as the associations of Bacteroides and Firmicutes in stool with obesity (Finucane et al. (2014)) and the associations of species in the vaginas of pregnant women with preterm birth (Callahan et al. (2017)).

The standard approach to countering taxonomic bias is to standardize the measurement protocol used within a given study. Statistical analyses are then conducted with the tacit assumption that all samples will be affected by bias in the same way and so the differences between samples will be unaffected. This argument is at least intuitively plausible for DA analyses based on multiplicative or fold differences (FDs) in a taxon’s abundance. If bias caused a species’ abundance to be consistently measured as 10-fold greater than its actual value, then we would still recover the correct FDs among samples. However, McLaren, Willis, and Callahan (2019) showed mathematically and with MGS measurements of artificially constructed (‘mock’) communities that consistent taxonomic bias can create fold errors (FEs) that vary across samples and, as a result, majorly distort cross-sample comparisons. In particular, they showed that the FE in a species’ proportion—the most common measure of relative abundance—varies among samples, distorting the observed FDs between samples. In some cases, bias can even lead to incorrect inferences about the direction of change (for example, by causing a taxon that decreased to appear to increase). Yet McLaren, Willis, and Callahan (2019) also found that other abundance measures—those based on the ratios among species—have constant FEs and may lead to more robust DA analyses. The implications of these findings for DA analysis of absolute abundances and for the joint analysis of variation of many species across many samples, as is typical in microbiome association testing, have yet to be investigated.

Here we use a combination of theoretical analysis, simulation, and re-analysis of published experiments to consider when and why taxonomic bias in MGS measurements leads to spurious results in DA analysis of relative and absolute abundances. We show that, in contrast to received wisdom, taxonomic bias can affect the results of DA methods that are based on species proportions, even if bias is the same across samples. We describe the theoretical conditions when this effect is negligible and when it may cause serious scientific errors, and explore this effect in case studies based on real microbiome experiments. We further demonstrate that another set of DA methods, based on the ratios among species, are robust to consistent bias. Finally, we present several methods for quantifying, correcting, or otherwise accounting for taxonomic bias in DA analyses which, in many cases, can be deployed with only modest changes to existing experimental and analytical workflows. These insights and methods will aid microbiome data analysts in turning the findings of microbiome studies into readily-translatable scientific knowledge.

## 2 How bias affects abundance measurements

This section extends the theoretical results of McLaren, Willis, and Callahan (2019) to describe how taxonomic bias in MGS experiments leads to errors in the relative and absolute abundances measured for various microbial species. All approaches to abundance quantification have systematic errors driven by taxonomic bias; however, some yield constant fold errors (FEs), while others yield FEs that depend on overall community composition and thus can vary across samples.

### 2.1 A model of MGS measurements

Our primary tool for understanding the impact of taxonomic bias on MGS measurement is the theoretical model of MGS measurement developed and empirically validated by McLaren, Willis, and Callahan (2019). This model describes the mathematical relationship between the read counts obtained by MGS and the (actual) abundances of the various species in a sample. Here we extend the model as first described McLaren, Willis, and Callahan (2019), which considers only relative abundances, to also consider absolute abundances. For concreteness, we will consider *absolute abundance*, or simply *abundance*, to refer to the number of cells per unit volume in a sample (cell concentration). That said, our results equally apply to other definitions of absolute abundance, such as the total number of cells in a sample or ecosystem and other abundance units such as biomass or genome copy number.

This model is the simplest that respects the multiplicative nature of taxonomic bias and the *compositional* nature of MGS measurements. The actual abundance of a species in a given sample, multiplied by its *measurement efficiency*—its rate of conversion from cells to taxonomically assigned sequencing reads—determines the species’ read count in that sample. Taxonomic bias presents as variation in the measurement efficiencies among species within an MGS experiment. The read counts further depend on sample-specific experimental factors that are typically unknown, such that they are best interpreted as only providing relative abundances (such data is said to be *compositional*; Gloor et al. (2017)).

We consider a set of microbiome samples measured by a specific MGS protocol that extracts, sequences, and taxonomically assigns reads to a set of microbial species *S*. We make several simplifying assumptions to facilitate our analysis and presentation. First, we consider only species-level assignment, and suppose that reads that cannot be uniquely assigned to a single species in *S* are discarded. Second, we ignore the possibility that reads are misassigned to the wrong species or the wrong sample. Third, we suppose that taxonomic bias acts consistently across samples at the species level—that is, a given species is always measured more efficiently than another to the same degree. Finally, unless otherwise stated, we treat sequencing measurements as deterministic, ignoring the ‘random’ variation in read counts that arise from the sampling of sequencing reads and other aspects of the MGS process. These assumptions, though unrealistic descriptions of most MGS experiments, serve the purpose of clearly demonstrating when and why consistent taxonomic bias creates errors in DA analysis.

Our model stipulates that the taxonomically-assigned read count for a species *i* in a sample *a* equals its abundance 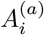 multiplied by a species-specific factor *B_i_* and a sample-specific factor *F*^(*a*)^,

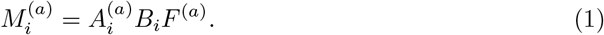

The species-specific factor, *B_i_*, is the *relative measurement efficiency* (or simply *efficiency*) of the species relative to an arbitrary baseline species (McLaren, Willis, and Callahan (2019)). The variation in efficiency among species corresponds to the taxonomic bias of the MGS protocol. The sample-specific factor, *F*^(*a*)^, describes the effective sequencing effort for that sample; it equals the number of reads per unit abundance that would be obtained for a species with an efficiency of 1. We can write the total number of assigned reads for the sample as

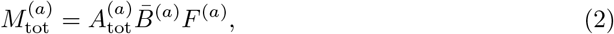

where 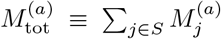 is the total read count and 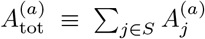 is the total abundance for all species *S*, and

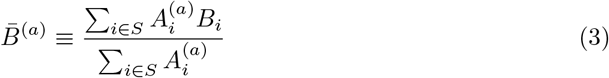

is the *sample mean efficiency*, defined as the mean efficiency of all species weighted by their abundance.

### 2.2 Relative abundance

We distinguish between two types of species-level *relative abundances* within a sample. The *proportion* 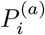 of species *i* in sample *a* equals its abundance divided by the total abundance of all species in *S*,

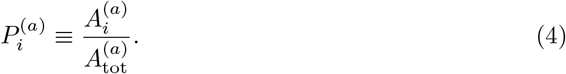

The *ratio* 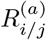 between two species *i* and *j* equals the abundance of *i* divided by that of *j*,

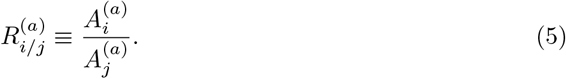

The measured proportion of a species is given by its proportion of all the assigned reads in a sample,

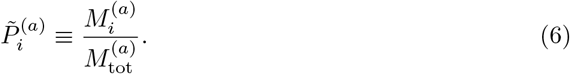

We use the tilde to distinguish the measurement from the actual quantity being measured. From Equations (1), (2), and (6), it follows that the measured and actual proportion are related by

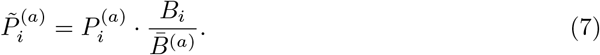

Taxonomic bias creates a fold-error (FE) in the measured proportion of a species that is equal to its efficiency divided by the mean efficiency in the sample. Since the mean efficiency varies across samples, so does the FE. This phenomenon can be seen for Species 3 in the two hypothetical communities in Figure 1. Species 3, which has an efficiency of 6, is under-measured in Sample 1 (FE < 1) but over-measured (FE > 1) in Sample 2. This difference occurs because the even distribution of species Sample 1 yields a mean efficiency of 8.33; in contrast, the lopsided distribution in Sample 2, which is dominated by the low-efficiency Species 1, has a mean efficiency of just 3.15. A demonstration in bacterial mock communities is shown in Figure 3C of McLaren, Willis, and Callahan (2019).

**Figure 1:**
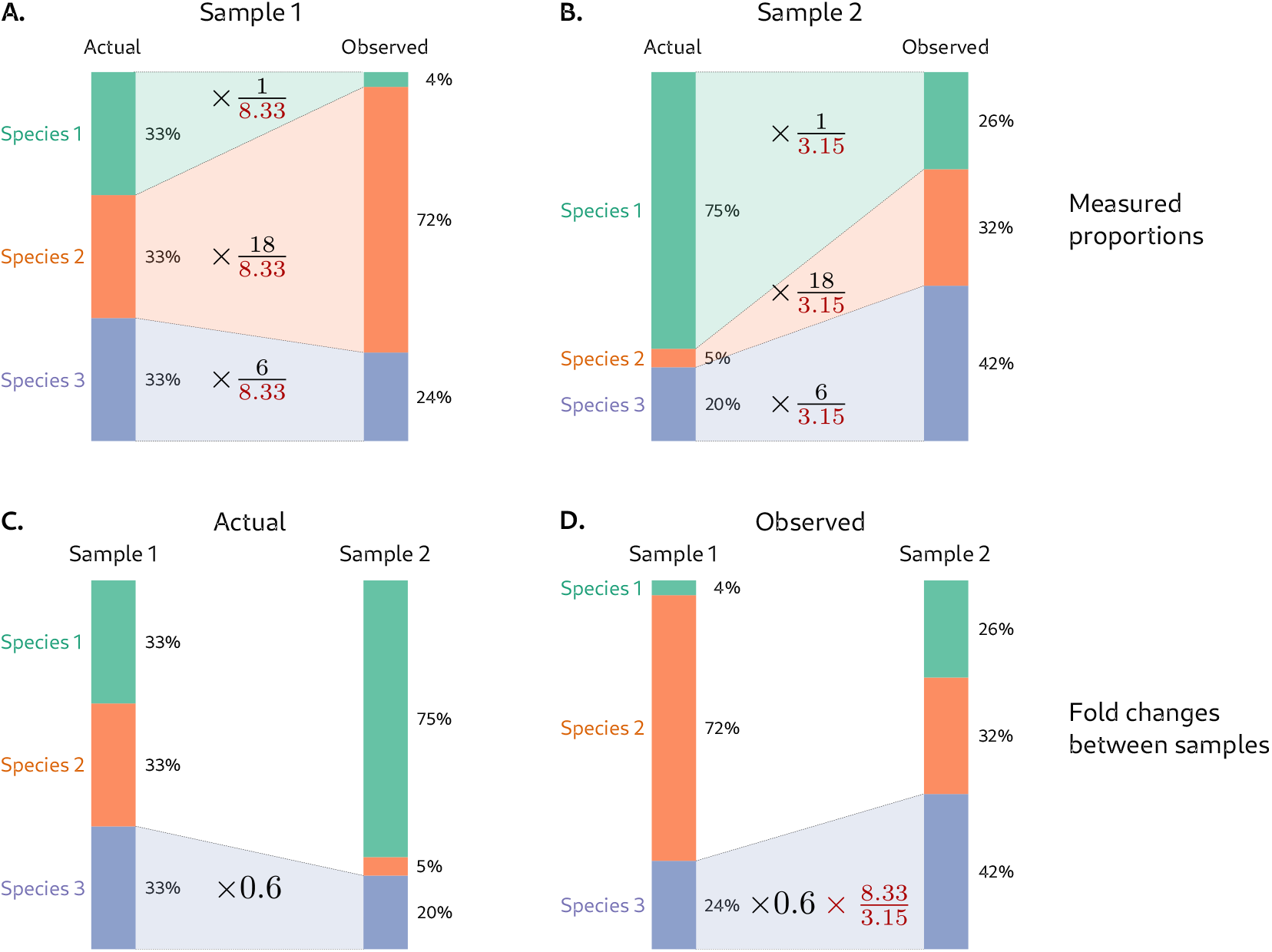
Taxonomic bias creates fold errors in species proportions that vary across samples and lead to inaccurate fold differences between samples. Top row: Error in proportions measured by MGS in two hypothetical microbiome samples that contain different relative abundances of three species. Bottom row: Error in the measured fold difference in the third species that is derived from these measurements. Species’ proportions may be measured as too high or too low depending on sample composition. For instance, Species 3 has an efficiency of 6 and is under-measured in Sample 1 (which has a mean efficiency of 8.33) but over-measured in Sample 2 (which has a mean efficiency of 3.15).

The measured ratio 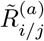 between species *i* and *j* is given by the ratio of their read counts,

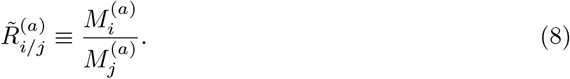

From Equations (1) and (8), it follows that the measured and actual ratio are related by

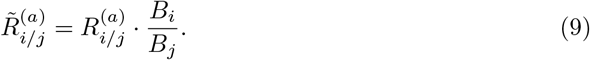

Taxonomic bias creates a FE in the measured ratio that is equal to the ratio in the species’ efficiencies; the FE is therefore constant across samples. For instance, in Figure 1, the ratio of Species 3 (with an efficiency of 6) to Species 1 (with an efficiency of 1) is over-measured by a factor of 6 in both communities despite their varying compositions. A demonstration in bacterial mock communities is shown in Figure 3D of McLaren, Willis, and Callahan (2019).

#### Higher-order taxa

We can consider a higher-order taxon *I*, such as a genus or phylum, as a set of species, {*i* ∈ *I*}. The abundance of taxon *I* in sample *a* is the sum of the abundances of its constituent species, 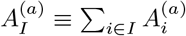. Similarly, the read count of taxon *I* is the sum 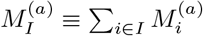. We further define the efficiency of taxon *I* as the abundance-weighted average of the efficiencies of its constituent species,

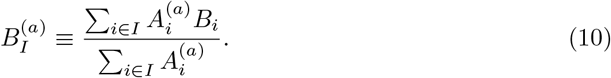

With these definitions, the read count for higher-order taxon *I* can be expressed as 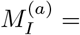 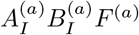. Thus 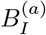 plays a role analogous to the efficiency of an individual species, but differs in that it is not constant across samples: If the constituent species have different efficiencies, then the efficiency of the higher-order taxon *I* depends on the relative abundances of its constituents and so will vary across samples (McLaren, Willis, and Callahan (2019)). As an example, suppose that Species 1 and Species 2 in Figure 1 were in the same phylum. The efficiency of the phylum would then be 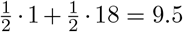 in Sample 1 and 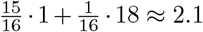 in Sample 2. Equations (7) and (9) continue to describe the measurement error in proportions and ratios involving higher-order taxa, so long as the sample-dependent, higher-order taxa efficiencies 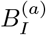 and 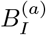 are used. In this way, we see that both proportions and ratios among higher-order taxa may have inconsistent FEs.

### 2.3 Absolute abundance

Several extensions of the standard MGS experiment make it possible to measure absolute species abundances. These extensions fall into two general approaches. The first approach leverages information about the abundance of the total community; for example, Vandeputte et al. (2017) measured total-community abundance using flow cytometry and multiplied this number by genus proportions measured by MGS to quantify the absolute abundances of individual genera (Vandeputte et al. (2017)). A second approach leverages information about the abundance of one or more individual species; for example, a researcher might ‘spike in’ a known, fixed amount of an extraneous species to all samples prior to MGS, and normalize the read counts of all species to the spike-in species (Harrison et al. (2021)). We consider each approach in detail to determine how taxonomic bias affects the resulting absolute-abundance measurements.

#### 2.3.1 Leveraging information about total-community abundance

Suppose that the total abundance of all species in the sample, 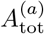, has been measured by a non-MGS method, yielding a measurement 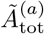. The absolute abundance of an individual species can be quantified by multiplying the species’ proportion from MGS by this total-abundance measurement,

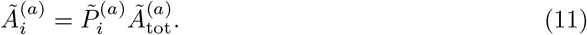

Total-abundance measurements recently used for this purpose include counting cells with microscopy (Lloyd et al. (2020)) or flow cytometry (Props et al. (2017), Vandeputte et al. (2017), Galazzo et al. (2020)), measuring the concentration of a marker-gene with qPCR or ddPCR (Zhang et al. (2017), Barlow, Bogatyrev, and Ismagilov (2020), Galazzo et al. (2020), Tettamanti Boshier et al. (2020)), and measuring bulk DNA concentration with a florescence-based DNA quantification method (Contijoch et al. (2019)).

Importantly, these methods of measuring total abundance are themselves subject to taxonomic bias that is analogous to, but quantitatively different from, the MGS relative abundance measurements. Flow cytometry may yield lower cell counts for species whose cells tend to clump together or are prone to lysis during steps involved in sample collection, storage, and preparation. Marker-gene concentrations measured by qPCR are affected by variation among species in extraction efficiency, marker-gene copy number, and PCR binding and amplification efficiency (Lloyd et al. (2013)). We can easily understand the impact of taxonomic bias on total-abundance measurement under simplifying assumptions analogous to those in our MGS model. Suppose that each species *i* has an *absolute efficiency* 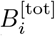 for the total-abundance measurement that is constant across samples. Further, let 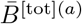 be the abundance-weighted average of these efficiencies in sample *a*—that is, the mean efficiency of the total-abundance measurement. Neglecting other error sources, the total-abundance measurement equals

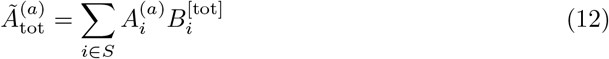

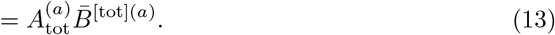

Species abundance measurements derived by this method (Equation (11)) are affected by taxonomic bias in both the MGS and total-abundance measurement. We can determine the resulting fold error (FE) in the estimate 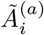 by substituting Equations (7) and (12) into Equation (11), yielding

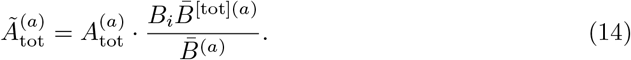

Equation (14) indicates that the FE in the measured absolute abundance of a species equals its MGS efficiency relative to the mean MGS efficiency in the sample, multiplied by the mean efficiency of the total measurement. As in the case of proportions (Equation (7)), the FE depends on sample composition through the two mean efficiency terms and so will, in general, vary across samples.

#### 2.3.2 Leveraging information about a reference species

Suppose that the absolute abundance of a *reference species r* has been fixed by the experimenter or been measured by independent means. This known or measured abundance 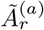 can be used in conjunction with the MGS read counts to obtain absolute abundances for all species. In the absence of taxonomic bias, the ratio of a species’ absolute abundance to its MGS read count is the same for all species in a given sample (Equation (1)). Hence the known ratio for the reference species can serve as conversion factor for obtaining the absolute abundance of a species *i* from its read count,

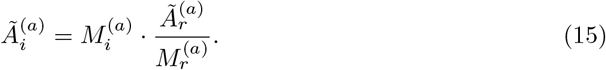

Let 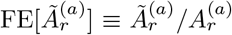 be the FE in the reference measurement. The effect on 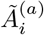 of taxonomic bias in the MGS measurement can be determined by substituting Equation (9) into Equation (15), yielding

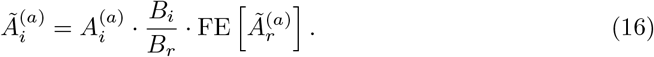

The FE in 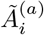 consists of two terms: the relative efficiency of species *i* to species *r* in the MGS measurement (*B_i_*/*B_r_*) and the FE in the reference species’ abundance 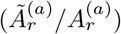.

A common application of this approach involves adding a ‘spike-in’ (as described above) in a known (and typically constant) abundance across samples (Stämmler et al. (2016), Ji et al. (2019), Tkacz, Hortala, and Poole (2018), Harrison et al. (2021), Rao et al. (2021)). In this case, the reference abundance 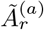 is determined from the concentration of the spike-in stock multiplied by the ratio of the spike-in to sample volumes.

Others have instead sought to determine naturally-occurring species that are thought to be constant across samples; we refer to such species as *housekeeping species* by analogy with the housekeeping genes used for absolute-abundance conversion in gene-expression studies (Silver et al. (2006)). Housekeeping species can sometimes be identified using prior scientific knowledge; for example, in shotgun sequencing experiments, researchers have used sequencing reads from the plant or animal host as a reference (Karasov et al. (2020), Regalado et al. (2020), Wallace et al. (2021)). A related approach involves computationally identifying species that are constant between pairs of samples (David et al. (2014)) or between sample conditions (Mandal et al. (2015), Kumar et al. (2018)). The abundance of a housekeeping species is typically unknown; therefore, to estimate the abundances of other species, we simply set 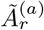 to 1 in Equation (15). The resulting abundance measurements have unknown but fixed units, which is sufficient for measuring fold changes across samples.

We suggest an additional way of using the reference-species strategy even in the absence of a spike-in or constant species: Performing targeted measurements of the absolute abundance of one or more naturally occurring species. These species can then be used as reference species in Equation (15) to measure the absolute abundances of all species. The most common form of targeted measurement involves using qPCR or ddPCR to measure the concentration of a marker-gene in the extracted DNA. It is also possible to directly measure cell concentration by performing ddPCR prior to DNA extraction (Morella et al. (2018)), flow cytometry with species-specific florescent probes, or CFU counting on selective media.

## 3 How bias affects DA results

We now turn to how the errors in MGS-based abundance measurements affect DA results. Our focus is on *multiplicative DA analyses* that estimate the fold differences (FD) in the relative or absolute abundance of species across time points or between experimental conditions. Multiplicative DA has a direct ecological interpretation via the multiplicative processes of exponential growth and decay. Many DA methods operate on additive differences in log-transformed abundances and are therefore multiplicative; these include popular methods such as DESeq2 (Love, Huber, and Anders (2014)), ALDEx2 (Fernandes et al. (2014)), corncob (Martin, Witten, and Willis (2020)), and time series methods based on the Lotka-Volterra model (e.g., Stein et al. (2013)). We also consider non-parametric rank-based methods; these methods are popular in case-control studies for which ecological interpretability is often of secondary importance to discovering stable associations between species and the condition of interest.

### 3.1 Fold change between a pair of samples

The building blocks of a multiplicative DA analysis are the measured FDs in a species’ abundance between pairs of individual samples. If the fold error (FE) in a species’ abundance measurement is constant across samples, then it will not affect the measured FDs between samples. In Section 2, we showed that consistent taxonomic bias creates a constant FE in the ratio between two species *i* and *j*, equal to the ratio in their efficiencies (Equation (9)). This error completely cancels when calculating the FD of this ratio from sample *a* to *b*,

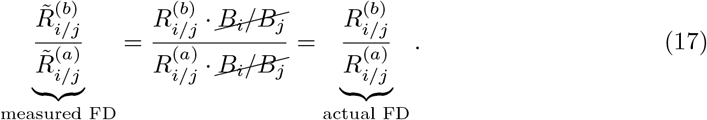

In contrast, bias creates a FE in the proportion of a species (Equation (7)) that varies inversely with the mean efficiency of the sample. Therefore, this error does not cancel when calculating the FD in the proportion,

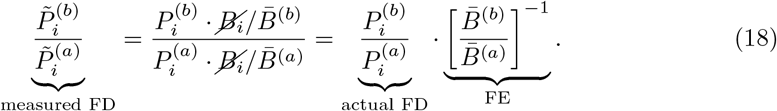

The varying mean efficiency does not cancel, leaving an FE in the FD of the species’ proportion that is equal to the inverse FD in the mean efficiency. Notably, this FE is the same for every species.

The different effects of taxonomic bias on the measured FDs in proportions versus ratios are illustrated in a hypothetical example in Figure 1 (bottom row). In this example, the mean efficiency decreases 2.6-fold from Sample 1 to Sample 2 (FD of 0.4), causing the FD in the proportion of each species to appear 2.6-fold larger than its true value. Though the FE in the FD is the same for each species, the biological implications of the error vary. In particular, there are three distinct types of error: an increase in the magnitude, a decrease in the magnitude, and a reversal in the direction of the measured FD. We can see each type of error in Figure 1. For Species 1, which increases in abundance and thus moves in the opposite direction of the mean efficiency, we see an increase in magnitude of the measured FD (actual FD: 2.3, measured FD: 6.5). For Species 2, which decreases and thus moves in the same direction as the mean efficiency but by a larger factor, we see a decrease in magnitude of the measured FD (actual FD: 0.15, measured FD: 0.44). For Species 3, which decreases but by a smaller factor than the mean efficiency, the species actually appears to increase—a reversal in direction of the measured FD (actual FD: 0.6, measured FD: 1.8). In contrast, the FDs in the ratios among species are identical in the actual and measured taxonomic profiles. For example, the ratio of Species 2 to Species 3 shows the same 4-fold decrease in the actual profiles (1 to 1/4) and in the measured profiles (3 to 3/4).

Taxonomic bias affects measured FDs in a species’ absolute abundance in a similar fashion. First, consider a species’ abundance measurement 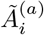 derived from a non-MGS measurement of total-community abundance using Equation (11). Equation (14) indicates that the FE in this measurement equals the (constant) species efficiency multiplied by 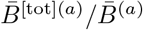, the ratio of the mean efficiency of the total-abundance measurement to that of the MGS measurement. This ratio can vary, creating error in the measured FD between samples. A notable special case is if the FDs in the mean efficiency of the total-abundance measurement mirrors that of the MGS measurement; then, the two will offset each other and lead to more stable FEs (and hence more accurate FDs) in 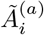. We discuss how this possibility might be exploited in real experimental workflows in Section 5.

Now consider a species’ abundance derived from a reference species using Equation (15). Equation (16) indicates that the FE equals a constant ratio in species’ efficiencies multiplied by the FE in the measurement of the reference species’ abundance. If the abundance of the reference species can be determined up to constant FE across samples, then the FE in 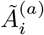 will also be constant, leading to accurate FDs.

### 3.2 Regression analysis of many samples

In most cases, a DA analysis can be understood as regression of microbial abundance variables against one or more covariates describing predictive properties of the sample. The simplest such regression relates a microbial abundance variable *y* to a covariate *x* and a *residual (or unexplained) error ε* via the simple linear regression formula

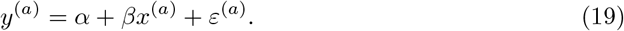

A multiplicative DA analysis can be conducted by setting *y* equal to the logarithm of the untransformed abundance measurement (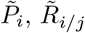, or 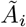). For example, *y* may be the log absolute abundance of species *i* 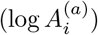 and *x* may be a continuous variable (such as pH) or a binary variable (such as *x* = 1 for treated patients and *x* = 0 for untreated controls). The regression coefficients *α* and *β* are parameters to be estimated from the data. A DA regression analysis is primarily interested in the slope coefficient *β*, which determines how the average or expected value of *y* increases with *x*. For a binary covariate, *β* equals the average difference in *y* between samples with *x* = 1 versus those with *x* = 0. When *y* equals the logarithm of a species’ relative or absolute abundance, the coefficient *β* equals the average log fold difference (LFD) in abundance between conditions corresponding to a unit increase in *x*. Although researchers often use more sophisticated models, the simple linear regression model in Equation (19) provides an intuitive basis for understanding the effect of taxonomic bias on regression-based DA analyses.

Appendix A describes a general framework for finding the error in the estimated slope coefficient 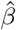 caused by taxonomic bias; here, we summarize and illustrate these results for the various abundance measures described in Section 2.

The effect that taxonomic bias has on the estimate 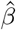 mirrors its effect on the measured FDs between a pair of individual samples: For abundance measures with a FE that is constant across samples, bias does not affect 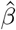. For abundance measures where the FE varies, bias can affect 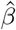, with its exact effect depending on how the mean efficiency varies across samples.

We illustrate the constant-FE case by considering a regression analysis of the log ratio between species *i* and *j*, 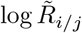. Taxonomic bias creates a constant FE in 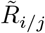 that translates to a constant additive error in 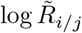. A constant (additive) shift in a response variable only affects the estimated intercept 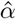 and not the estimated slope 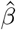. Thus, in this case, taxonomic bias does not affect 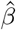.

We illustrate the varying-FE case by considering the absolute abundance of a species *i* obtained by multiplying the species’ MGS proportion by a non-MGS measure of total-abundance, using Equation (11). For simplicity, suppose that total-community abundance has been measured completely accurately. In this case, the measured abundance 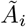 has a FE that varies inversely with the mean efficiency 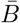. If 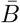 is stable across samples, then the additive error in 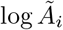 will be approximately constant and 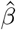 is again unaffected—bias has no effect on the DA result. If, on the other hand, 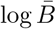 tends to linearly vary with the covariate *x*, then the additive error in 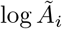 will linearly vary in the opposite direction and cause a systematic error in 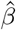 that is equal in magnitude but opposite in sign to the slope of 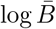 with *x*. A third possibility is that 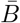 varies in an essentially random manner that is independent of the covariate *x*. In this scenario, the error from taxonomic bias in the measurement 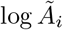 acts to increase the noise in 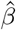 (i.e., increase its standard error), but does not cause a systematic error.

Figure 2 shows a simulated example of a regression analysis of the absolute abundances of 10 species across two conditions in which the mean efficiency is systematically greater in the condition *x* = 1 than in the condition *x* = 0. The increase in the log mean efficiency from *x* = 0 to *x* = 1 (Figure 2 A and B) corresponds to an artificial decrease in the estimated LFD between conditions for each species (Figure 2 C and D). The absolute error created by bias on the LFD estimate is the same for each species; however, its scientific impact varies. For the three species with large positive LFDs (Species 9, 10, and 1), bias decreases the magnitude of the LFD estimate. In contrast, for the three species with large negative LFDs (Species 8, 2, and 3), bias increases the magnitude of the estimate. For one species with a small positive LFD (Species 5), the effect of bias results in a negative LFD estimate (direction error). The three remaining species (Species 7, 6, and 4) have LFDs near zero, but bias causes them to have large negative LFDs.

**Figure 2:**
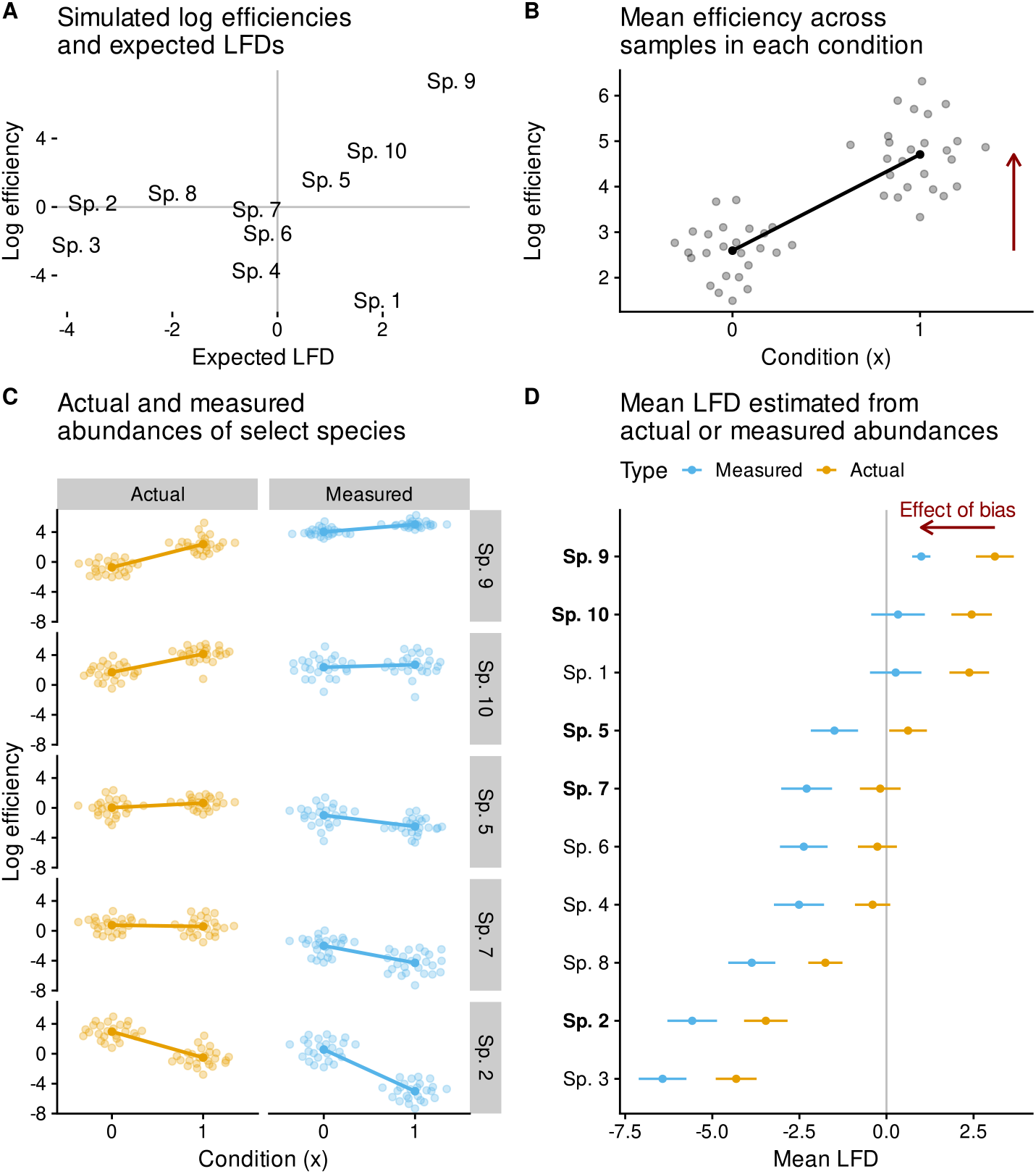
Taxonomic bias distorts multi-sample differential abundance analysis when the mean efficiency of samples is associated with the covariate of interest. This figure shows the results of a regression analysis of simulated microbiomes consisting of 10 species and 50 samples from two environmental conditions, which are indexed by *x* = 0 and *x* = 1. Panel A shows the (log) measurement efficiencies of the 10 species against their differential-abundance parameter values (expected LFD in absolute abundance between a sample from condition *x* = 1 to a sample from condition *x* = 0). The simulation was created so that the species with the largest efficiency (Sp. 9) also has the largest positive expected LFD. The increased relative abundance of Sp. 9 in condition *x* = 1 drives an increase in the (log) mean efficiency (Panel B), which induces a systematic and consistent negative shift in the estimates of the mean LFD for each species (Panels C and D). Panel D shows the estimated mean LFD with 95% confidence interval for each species, when either estimated the actual abundances or the inaccurate measured abundances. The error (estimate from measured abundances minus estimate from actual abundances; red arrow in Panel D) equals the negative mean LFD the mean efficiency (red arrow in Panel B).

### 3.3 Rank-based analyses

Rank-based DA methods work by first applying a cross-sample rank-transformation to the abundance variable *y*: The sample in which *y* has the smallest value receives a rank of 1, the sample with the second smallest value receives a rank of 2, etc. These methods then analyze how the average rank of *y* varies across sample conditions (Conover (2012)). Rank-based methods commonly used in microbiome DA analysis include the Wilcoxon-Mann-Whitney and Kruskal-Wallis tests for a discrete covariate (e.g. case vs control) and Spearman’s rank correlation for a continuous covariate (e.g. pH); these tests have collectively been applied to species proportions (Callahan et al. (2017),Fettweis et al. (2019)), ratios (Fernandes et al. (2014)), and absolute abundances (Vieira-Silva et al. (2019)).

Multiplying a variable by a constant factor does not change its cross-sample ranks, making rank-based methods unaffected by a constant FE in the abundance variable *y*. Thus in our MGS measurement model, taxonomic bias does not affect rank-based DA analyses of species ratios or absolute abundances derived from a reference species. But for species proportions and absolute abundances derived from a total-community abundance measurement, variation in the mean efficiency 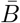 can affect the cross-sample ranks and cause error in DA results if 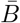 is associated with the covariate. Further work is needed to determine whether rank-based DA methods are significantly more robust to variation in 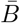 than multiplicative DA methods.

## 4 Case studies

To better understand the potential impact of taxonomic bias on DA analysis in practice, we conducted several case studies spanning a range of biological scenarios and sequencing technologies.

### 4.1 Foliar fungi experiment

The taxonomic bias of species in ‘mock communities’ of known composition can be directly measured, and the measured bias can then be used to correct downstream DA analysis (McLaren, Willis, and Callahan (2019)). In practice it is difficult to construct control communities that span the many species present in complex natural communities. However, gnotobiotic community experiments are well suited to this form of *calibration via community controls* since it is possible to assemble mock communities containing all species in known relative abundances.

In a study of host-commensal-pathogen interactions, Leopold and Busby (2020) inoculated plants with 8 commensal fungal species and subsequently exposed plants to a fungal pathogen. The authors used ITS amplicon sequencing to measure communities before and after pathogen infection. Motivated by the substantial ribosomal copy-number variation (CNV) in fungi (Lofgren et al. (2019)), the authors also performed control measurements of mock communities that they constructed from quantified genomic DNA of the 9 species in the experiment; these controls were used to measure taxonomic bias with the method of McLaren, Willis, and Callahan (2019). The authors found a 13-fold difference between the most and least efficiently measured commensal, while the pathogen was measured 40-fold more efficiently than the least efficiently measured commensal.

Leopold and Busby (2020) performed two related DA analyses on the pre-infection communities: the first characterized the relative importance of host genetics and species arrival order on species relative abundances in the fully-established community, and the second quantified the strength of ‘priority effects’—the advantage gained by a species from being allowed to colonize first. Both analyses were based on fold differences (FDs) in species proportions and so in principle were sensitive to taxonomic bias. To improve accuracy, the authors incorporated the bias measured from the control samples into analysis-specific calibration procedures.

We repeated the two DA analyses of Leopold and Busby (2020) with and without calibration and found that the results did not meaningfully differ (SI Analysis of host genetics and arrival order, SI Analysis of priority effects). To understand why, we examined the variation in species proportions and the mean efficiency across the pre-infection communities (SI Analysis, SI Figure 5). Despite the 13-fold variation in the efficiencies among species, the mean efficiency hardly varied across samples (SI Figure 5C), having a geometric range of 1.62 and a geometric standard deviation of 1.05. This consistency in the mean efficiency was despite the fact that each species showed substantial multiplicative variation (SI Figure 5A). But the pre-infection samples were always dominated by the three species with the highest efficiencies, which varied by 3-fold and by just 1.5-fold between the two most dominant. The mean efficiency, equal to the proportion-weighted arithmetic average of species efficiencies, is insensitive to species present at low relative abundance and so remained relatively constant across samples. Because the multiplicative variation in the mean efficiency was much smaller than that in the proportions of individual species, it had a negligible impact on the inferred FDs and the DA analyses based on them.

We performed an additional DA analysis on the data from Leopold and Busby (2020) to investigate whether any commensals increased in absolute concentration in response to infection (SI Analysis). The pathogen is absent in pre-infection samples but tends to dominate the community post-infection, resulting in a substantially higher mean efficiency in post-infection samples (SI Figure 6). Across different host genotypes, the average post-infection increase in mean efficiency ranged from 2.5-fold to 5.2-fold. Using gamma-Poisson regression of the observed read counts, we estimated the average LFD in commensal species proportions following infection with and without calibration (SI Figure 7). The commensals’ proportions typically decreased post-infection, which is expected given the pathogen’s growth to most abundant community member and the sum-to-one constraint faced by proportions. However, failure to account for bias caused the decrease to be overestimated, by an amount corresponding to the inverse change in log mean efficiency. For commensal-host pairs with relatively small observed decreases, bias correction greatly reduced the magnitude of the negative LFDs or, in several cases, resulted in LFDs that were near 0 or slightly positive.

Although Leopold and Busby (2020) did not include absolute-abundance measurements, we can consider the impact taxonomic bias would have on an absolute DA analysis in a simple scenario in which total genome concentration of each pre- and post-infection sample is perfectly known. We consider the approach of multiplying the total genomic concentrations of each sample with the species proportions measured by MGS as described by Equation (11). In this case, the bias in the MGS measurements will create absolute errors in the estimated LFDs of genome concentration of equal magnitude to the errors in the LFD estimates for proportions. If total abundances also differed after pathogen growth, however, we may make directional errors in determining the FD. Suppose that total abundance increased by 2-fold due to pathogen growth; in this case, species whose proportions remained approximately constant would have increased in absolute abundance by around 2-fold. Bias shifts FD estimates downwards by 2.5-fold to 5.2-fold (depending on host genotype). Hence, without bias correction, we would instead conclude that these species decreased.

### 4.2 Vaginal microbiomes of pregnant women

A growing number of MGS studies have used DA analysis to describe associations between members of the vaginal microbiome and health conditions including urinary tract infections, sexually-transmitted infections, bacterial vaginosis, and preterm birth. However, these associations often vary between studies. DA analyses of the vaginal microbiome are commonly based on proportions, creating an opportunity for taxonomic bias to impact results. Substantial taxonomic bias has been experimentally demonstrated in MGS protocols applied to vaginal samples or in vitro samples of vaginally-associated species (Yuan et al. (2012), Brooks et al. (2015), Gill et al. (2016), Graspeuntner et al. (2018)). The different biases of different MGS protocols have been proposed as one explanation for discrepancies in DA results across studies (Callahan et al. (2017)).

As part of the Multi-Omic Microbiome Study: Pregnancy Initiative (MOMS-PI) study, Fettweis et al. (2019) collected longitudinal samples from over 1500 pregnant women, including nearly 600 that were taxonomically profiled by amplicon sequencing of the 16S V1-V3 hypervariable region. The taxonomic bias of the MOMS-PI MGS protocol was previously quantified by McLaren, Willis, and Callahan (2019) using control measurements by Brooks et al. (2015) of cellular mock communities of seven common, clinically-relevant vaginal bacterial species. Of these, *Lactobacillus iners* had the highest efficiency, which was nearly 30-fold larger than that of the species with the lowest efficiency, *Gardnerella vaginalis*. A second *Lactobacillus* species, *L. crispatus*, had an efficiency that was approximately 2-fold less than *L. iners* and 15-fold greater than *G. vaginalis*. These species, along with the unculturable *Lachnospiraceae BVAB1*, are most frequently the most abundant species in a sample and commonly reach high proportions of over 70% (SI Analysis). Therefore, shifts between these dominant species could drive large changes in the sample mean efficiency, which might in turn distort DA analyses.

We sought to understand the potential impact of bias on DA analysis of vaginal microbiome profiles from the MOMS-PI study. We first examined variation in the mean efficiency across samples under the assumption that bias in the MOMS-PI study was accurately represented by the Brooks et al. (2015) control measurements. Using taxonomic relatedness to impute the efficiencies of species not in the controls, we were able to calibrate the MOMS-PI profiles and estimate the mean efficiency of each sample (SI Analysis). The mean efficiency varies substantially across vaginal samples (Figure 3), with samples in which a *Lactobacillus* species is most abundant typically having a mean efficiency that is 3-fold to 20-fold greater than samples in which *G. vaginalis* is most abundant. The vaginal microbiome sometimes shifted between *Lactobacillus*-dominance and *Gardnerella*-dominance between consecutive visits in individual women (SI Figure 8), which caused magnitude and direction errors in the trajectories of lower-abundance species (SI Figure 9).

**Figure 3:**
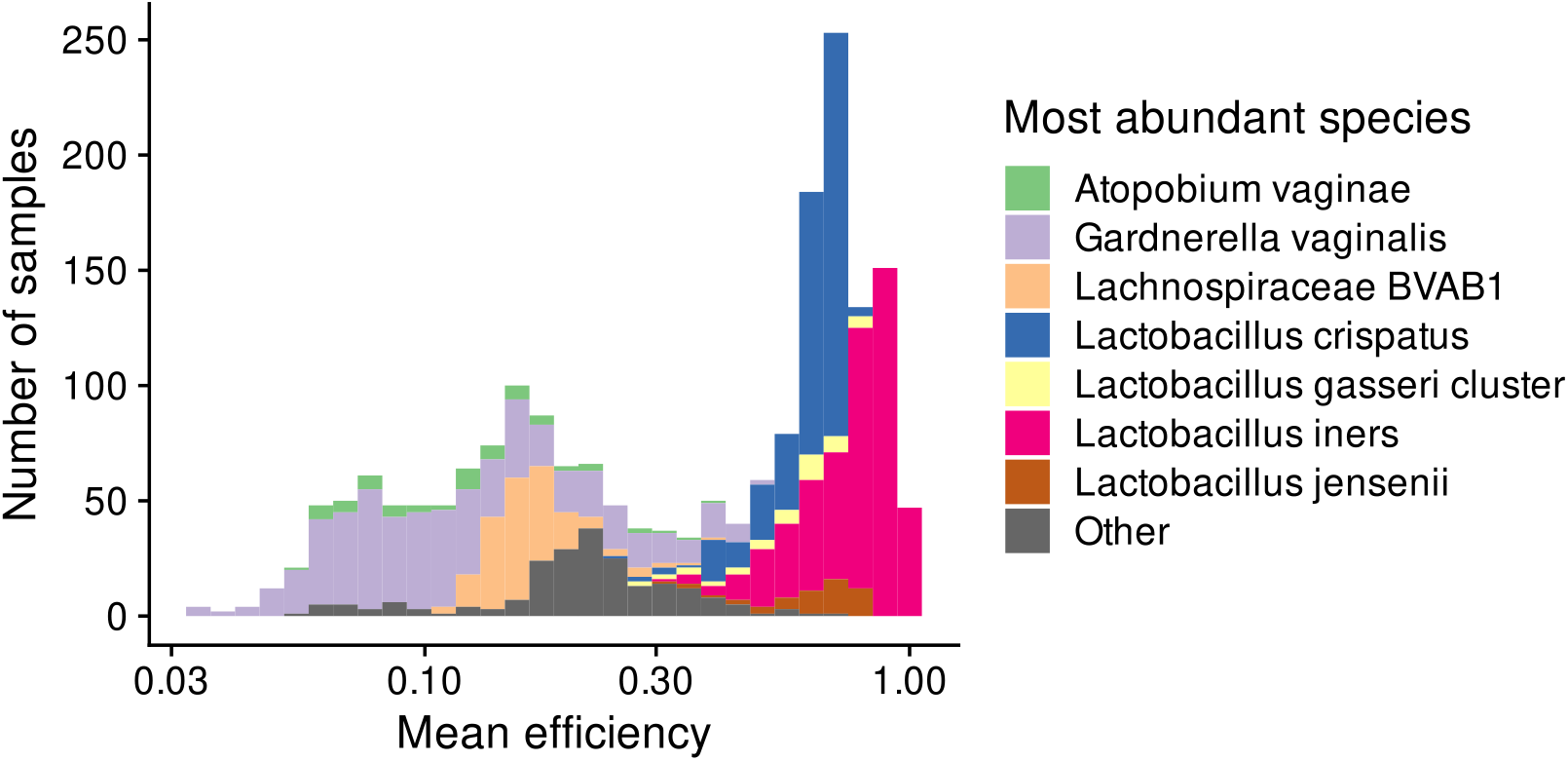
The mean efficiency in vaginal samples from the MOMS-PI study varies with the most abundant species.

These observations suggest that a proportion-based DA analysis of a health condition that is associated with *Lactobacillus* and/or *Gardnerella* would be prone to spurious results. To demonstrate this possibility, we performed a DA analysis of the MOMS-PI samples with respect to a simulated health condition (SI Analysis). Bacterial vaginosis (BV) has been repeatedly found to be associated with a dramatically reduced proportion of *Lactobacillus spp*., an increased proportion of *G. vaginalis*, and higher within-community (alpha) diversity (Srinivasan et al. (2012), Cartwright et al. (2018)). As a proxy for clinical BV, we split samples into high diversity (BV-like), intermediate diversity, and low diversity (non-BV-like) groups based on Shannon diversity in observed (uncalibrated) microbiome profiles. The mean efficiency was lower in the high-diversity BV-like samples (mean LFD of −1.6) due to a lower abundance of *Lactobacillus spp*.. We estimated the LFD in proportion from non-BV-like to BV-like samples for 30 prevalent bacterial species, with and without bias correction, using gamma-Poisson (or negative-binomial) regression of the observed read counts to fit a linear model to log species proportions. We accounted for bias by including an offset term in the linear model equal to the log ratio of the species’ efficiency to the sample mean efficiency. As expected, bias correction reduced the estimated LFD from non-BV-like to BV-like groups for every species tested, though the importance of this effect varied among species (SI Figure 10).

### 4.3 Human gut microbiomes

We wondered whether DA studies of the human gut microbiome might be less affected by bias than studies of the vaginal microbiome, due to the greater alpha diversity observed within gut microbiome samples. Recall that the mean efficiency is an abundance-weighted average over the species in a sample. Thus, all else equal, we expect the mean efficiency to vary less across samples from ecosystems where samples have higher alpha (within-sample) diversity, as is the case for stool samples relative to vaginal samples (Huttenhower et al. (2012)). On the other hand, the proportion of individual species may also vary less less across samples when alpha diversity is higher, in which case the effect of bias on DA results need not diminish.

To investigate this question, we analyzed bacterial species profiles of vaginal and stool samples derived from shotgun sequencing in the Human Microbiome Project (Huttenhower et al. (2012), SI Analysis). On average, stool profiles had substantially greater order-2 alpha diversity (Inverse Simpson index) than vaginal profiles (SI Figure 11A; geometric mean (GM) ×/ geometric standard deviation (GSD) of 6.3 ×/ 1.7 in stool samples and 1.4 ×/ 1.6 in vaginal samples). To assess the potential importance of bias for proportion-based DA analyses in the two ecosystems, we quantified the multiplicative variation in the mean efficiency across samples for a large number of possible taxonomic biases under the assumption that the measured profiles reflected the truth. Across simulation replicates, the GSD in the mean efficiency was typically lower in stool than in vaginal samples (SI Figure 11B; ratio of GSD in vaginal samples to GSD in stool samples of 1.6 ×/ 1.5 across 1000 simulations). Notably, however, the multiplicative variation in species also tended to be lower (SI Figure 11C); the GSD in the proportion of a gut species across gut samples was an average of 1.75-fold lower than that of a vaginal species across vaginal samples. Recall that the importance of bias estimation of the FD in a species’ proportion depends on GSD in the mean efficiency relative the GSD in the species’ proportion (Section 3). Thus these results suggest that although the mean efficiency likely varies less across stool than vaginal samples, bias may be just as problematic for proportion-based DA analyses.

### 4.4 Microbial growth in marine sediments

The surface layers of marine sediments harbor diverse and abundant microbiomes. Total cell density and species richness decrease with depth as resources are consumed in older, deeper sediment layers; however, some taxa are able to grow and increase in density as they are slowly buried. Lloyd et al. (2020) performed a systematic assessment of growth rates of bacterial and archaeal taxa over a depth of 10 cm (corresponding to ~40 years of burial time) in sediment of the White Oak River estuary. Taxa proportions were measured with 16S amplicon sequencing and total community densities were measured by counting cells using epifluorescence microscopy. The authors then calculated cell concentrations for each taxon by multiplying the taxon’s proportion by the total concentration, a method they referred to as FRAxC measurements for ‘fraction of 16S reads times total cell counts’. Taxon-specific growth rates were inferred from the slope of a simple linear regression of the log of the calculated concentration of each taxon against burial time over the first 3 cm below the bioirrigation layer (corresponding to ~8 years of burial).

The FRAxC concentration measurements made by Lloyd et al. (2020) correspond to the total-abundance approach to inferring absolute abundance (Equation (11)). Accordingly, taxonomic bias could lead to systematic error in FRAxC-derived growth rates if sample mean efficiency tends to systematically vary with burial time. Such systematic variation could occur if microbes with tougher cell walls both persist longer in the sediment and are more difficult to extract DNA from than microbes with weaker cell walls. In this scenario, the relative abundance of tougher species will increase with burial time, as the cells of weaker species degrade. This shifting composition will cause the sample mean efficiency to decrease with time, leading to inflated growth-rate estimates for all taxa. Taxa that decay with depth sufficiently slowly would mistakenly be inferred to have positive growth from the FRAxC-calculated abundances.

Importantly, Lloyd et al. (2020) also used qPCR to measure the absolute abundance of two taxa from these same samples. By comparing qPCR to FRAxC growth rates for these taxa, we can estimate the systematic error in FRAxC growth rates (SI Analysis) Because the systematic error that bias causes in the regression slope is the same for each taxon (Section 3), these comparisons allow us to draw conclusions about the accuracy of FRAxC growth rates for all taxa.

The first soil core included qPCR measurements of a single archaeal taxon, *Bathyarchaeota*, for which growth rates by qPCR and FRAxC were nearly identical (doubling rates of 0.099/yr by FRAxC and 0.097/yr by qPCR). The second soil core included qPCR measurements of *Bathyarchaeota* and a second taxon, *Thermoprofundales*/MBG-D. In this core, FRAxC and qPCR growth rates differed more substantially, with growth rates from FRAxC being larger by 0.012/yr for *Bathyarchaeota* (0.112/yr by FRAxC and 0.1/yr by qPCR) and by 0.086/yr for *Thermoprofundales*/MBG-D (0.294/yr by FRAxC and 0.208/yr by qPCR). Uncertainty in the FRAxC- and qPCR-derived growth rates is not reported and is likely substantial; however, the fact that the FRAxC-derived rates are larger than qPCR-derived rates in all three cases is consistent with our hypothesis that mean efficiency decreased with depth in a manner that systematically biased FRAxC-derived rates to higher values. The differences in growth rate are small in absolute terms; however, the maximum observed difference of 0.086/yr suggests an error large enough to impact results for some taxa classified as positive growers, whose FRAxC growth rates ranged between 0.04/yr and 0.5/yr. In summary, our comparison between FRAxC and qPCR measurements supports the overall study conclusion that many taxa did grow following sediment burial; however, we should remain uncertain as to whether species with small, positive FRAxC-derived growth rates were in fact growing or rather were slowly declining in abundance.

### 4.5 Summary and discussion

The effect that consistent taxonomic bias has on proportion-based DA analyses depends on the MGS protocol, the biological system, and the sample comparisons under investigation. Although our case studies explore a limited range of possibilities, some general patterns stand out.

In several cases, the mean efficiency remained stable across samples, so that LFD esimates were unaffected by bias. In the pre-infection fungal microbiome samples of Leopold and Busby (2020), we observed that the mean efficiency remained stable despite substantial bias and large multiplicative variation in species proportions among samples. Because the variation in the mean efficiency was much less than that of the individual species, the impact of bias on DA analysis was negligible. In the vaginal case study, we also observed that the mean efficiency was relative stable across vaginal microbiomes that were dominated by the same species, despite substantial bias and large LFDs in non-dominant species. In both cases, the stability of the mean efficiency can be understood by the fact that it is an additive average over species and so is primarily determined by the most abundant species in the sample.

Yet we also observed cases where the mean efficiency varied substantially. In several cases, the (log) mean efficiency co-varied sufficiently strongly with the condition of interest to cause substantial systematic errors in LFD estimates. Examples include the comparison of foliar fungal microbiomes pre- and post-infection, for which the mean efficiency substantially increased post-infection, and the comparison of vaginal microbiomes with low versus high diversity in the MOMS-PI study, for which the mean efficiency was typically lower in high-diversity samples. Our analysis of marine sediment communities is consistent with a systematic decline of the mean efficiency with burial time that is sufficient to substantially inflate the estimated growth rates of slowly changing species. The FD in the mean efficiency is bounded by the largest FD of any species’ proportion. Therefore, the error in the estimated LFDs tend to be practically significant only for species whose LFDs are substantially smaller in magnitude than the largest LFD.

Variation in the mean efficiency may also be substantial, yet unassociated with the condition of interest. Although we did not directly observe this scenario directly, we have reason to think that it may be common in real microbiome studies. Our simulation analysis of gut microbiomes suggest that even in diverse ecosystems, bias will often cause multiplicative variation in the mean efficiency that is comparable to that of individual species. This variation is much less problematic for DA results when it not associated with the condition under study, since any loss in precision can in principle be offset by an increase in sample size. For certain applications, however, it will be important to remember that the LFDs between individual samples remain unreliable.

Under what conditions should we expect the problematic third scenario? Our case studies suggest one prominent mechanism for causing systematic variation in the mean efficiency: the existence of one or more species with unusually high or low efficiencies that tend to dominate the community more often in one sample condition than another. This mechanism was responsible for the negative effect of bias in our DA analyses of foliar fungal microbiomes following infection and of vaginal microbiomes with low versus high diversity. In experimental systems where no one species frequently forms a large fraction of the community, systematic variation of the mean efficiency can still arise through the collective change of many species that are associated with the condition of interest. We described a plausible instance in the marine-sediment case study, where increased burial time selects for lysis-resistant (and so lower efficiency) species. As another example, treatment with an antibiotic might selectively kill Gram negative species, which also tend to be easier to lyse, thereby decreasing the mean efficiency in fecal samples collected after treatment. A generic mechanism which might spawn such condition-efficiency associations stems from the evolutionary relationships among species. Species with recent common ancestry are expected to share a number of traits that determine measurement efficiency, including cell wall structure, genome size, ribosomal copy number, and PCR binding sequence. They are also expected to share traits related to the condition of interest. If related species tend to have similar efficiencies and similar associations with the condition, then they may drive an association of the mean efficiency with the condition.

These observations provide reasons to both worry and hope. It seems likely that in many studies, the mean efficiency is consistent (first scenario) or is at least not associated with the condition (second scenario), so LFD estimates remain accurate (or at least not overconfident). Yet it is not obvious which scenario any study falls into. There are plausible mechanisms leading to systematic variation of the mean efficiency even in ecosystems with high species diversity. Moreover, our explorations suggest that random variation in the mean efficiency is common and distorts comparisons between individual samples. Thus while we should not discount the large set of existing DA results, we should seek ways to better assess the robustness of results from previous studies and to measure and correct the error caused by bias in future ones.

## 5 Solutions

### 5.1 Ratio-based relative DA analysis

Since bias creates constant fold errors (FEs) in the ratios among species, it can be countered by using relative DA methods that analyze fold differences (FDs) in these ratios. A variety of ratio-based methods have been developed under the framework of Compositional Data Analysis (CoDA; Gloor et al. (2017)). CoDA methods may more generally consider the ratios between products of multiple species, possibly raised to some exponent; a simple example is the geometric mean of several species. The mathematical operations of multiplication and exponentiation maintain constant FEs. Hence such products provide a method for aggregating species into a higher-level taxonomic units that maintains bias invariance when estimating FDs. Ratio-based analyses also avoid the potential for misinterpreting differences in a species’ proportion that are driven by the sum-to-one constraint (Gloor et al. (2017)).

We note two important caveats. First, in most MGS datasets, any given species is likely to be extremely rare in most samples, such that no sequencing reads are observed. In order to analyze FDs, DA methods interpret these zero counts as evidence that the true abundance of the species is very small (but still positive). A common approach to enable the computation of FDs in the presence of zeros is to add a small value, or ‘pseudo-count’, to the read counts, making all values positive. Unfortunately, this procedure violates bias invariance. How bias interacts with more sophisticated approaches for handling zeros remains an important open question. Second, the bias invariance of ratios among species does not extend to additive aggregates of species into higher-order taxa such phyla unless bias is conserved within the species group (McLaren, Willis, and Callahan (2019)). Multiplicative aggregates of species, recently used for biomarker discovery (Rivera-Pinto et al. (2018), Quinn and Erb (2020)), provide an alternative that remains bias-invariant but are harder to interpret.

If biological or technical considerations favor a DA method that is not bias invariant, it can be beneficial to also use a ratio-based method as a robustness check. For example, Hevroni et al. (2020) examined how the within-sample ranks of viral species changed from summer to winter. Since changes in within-sample ranks can be affected by bias, the authors also considered the changes in the centered log ratios (CLR) associated with each species. They found that the species whose within-sample ranks increased also increased in their CLR value, providing evidence that their initial results were not driven by bias.

### 5.2 Calibration using community controls

*Community calibration controls* are samples whose species identities and relative abundances are known, either by construction or by characterization with a chosen ‘gold standard’ or *reference protocol* (McLaren, Willis, and Callahan (2019), Clausen and Willis (2022)). Inclusion of these samples along with the primary samples in an MGS experiment can enable researchers to calibrate their MGS measurement by directly measuring and removing the effect of bias. Calibration can be extended to species not in the controls by imputing the efficiencies of the missing species using phylogenetic relatedness, genetic characteristics (such as 16S copy number), and/or phenotypic properties such as cell-wall structure.

Calibration makes it possible to correct for bias in relative and absolute DA methods that would otherwise not be bias invariant. To demonstrate, we used calibration to improve the estimated FDs between samples in the absolute abundance of a species in the bacterial mixtures from Brooks et al. (2015). We used the bias estimated from a single 7-species mixture to calibrate the MGS-derived proportions in all measured mixtures. We obtained calibrated absolute abundances by multiplying the calibrated proportions by the known total abundance (Equation (11)). Calibration greatly increased the accuracy of the estimated FDs (Figure 4, A vs. B). The fungal and vaginal case studies of Section 4 provide several practical examples of calibration in proportion-based DA analyses.

**Figure 4:**
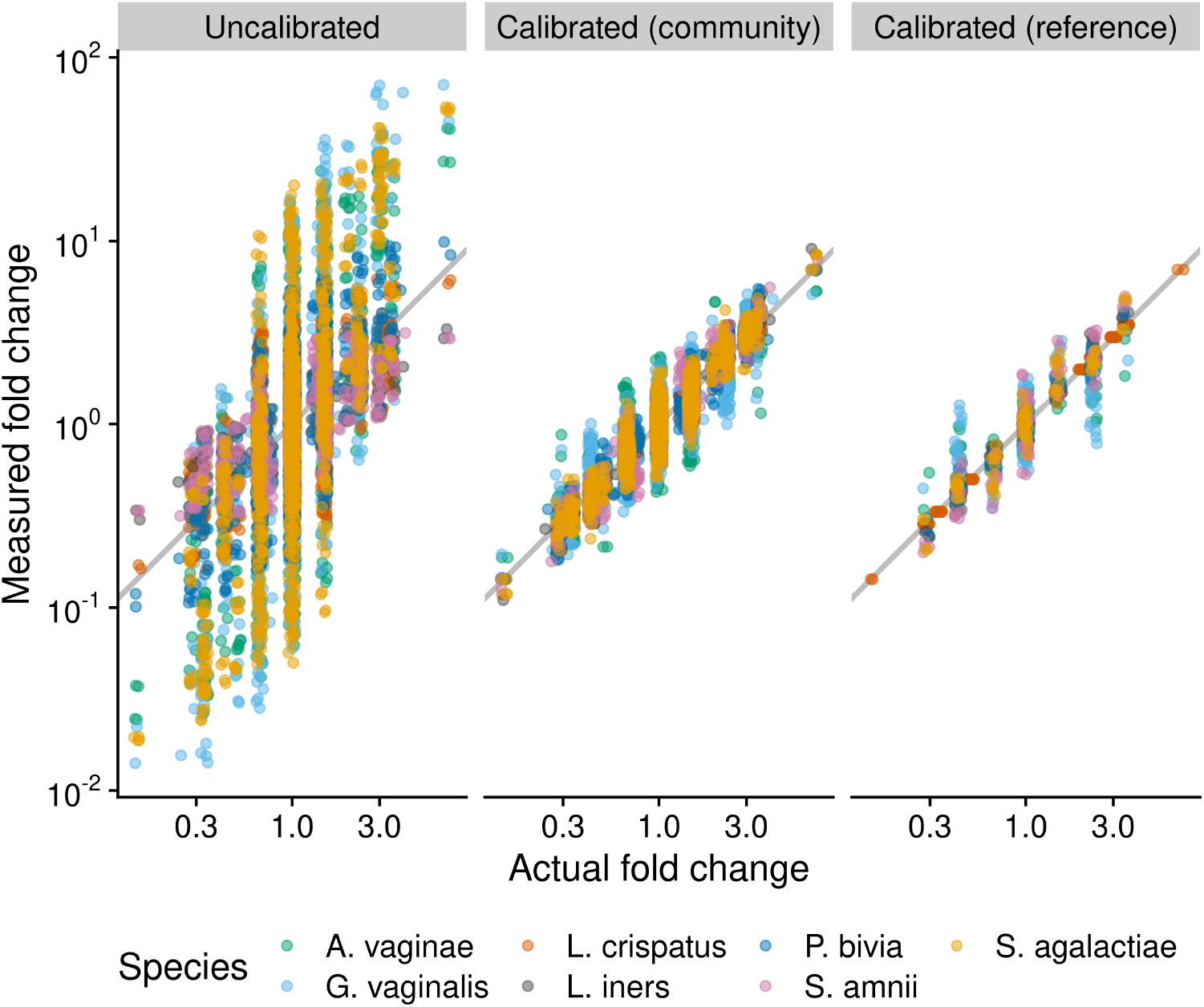
Fold differences can be calibrated using community controls or reference species. The figure compares the performance of three methods for measuring fold differences (FDs) in absolute cell density in cellular mock communities of 7 vaginal species, which were constructed and measured via 16S sequencing by Brooks et al. (2015). The ‘Uncalibrated’ FDs are derived directly from uncalibrated individual abundance measurements, which equal the product of the species’ proportion by the total density (which is constant by construction). The ‘Calibrated (community)’ measurements are computed from abundance measurements where the proportions are first corrected for the taxonomic bias that was estimated from a single sample that contained all 7 species. The ‘Calibrated (reference)’ measurements are computed from abundances measured with the reference-species method, with *Lactobacillus crispatus* used as the reference; that is, the true abundance of *L. crispatus* is treated as known and used to infer the abundance of the remaining 6 species. Only samples that contain *L. crispatus* are included.

In studies of synthetic communities (like the fungal case study) or of ecosystems dominated by a relatively small number of culturable taxa (like the vaginal microbiome), calibration controls can be artificially constructed (‘mock’) communities. In other cases, *natural community controls* can be derived from aliquots of a homogenized microbiome sample. Measurement of these controls across multiple experiments enables characterization of the *differential bias* between experiments (McLaren, Willis, and Callahan (2019)). Calibration using differential bias can make results directly comparable across studies that used different protocols.

### 5.3 Absolute-abundance methods with more stable FEs

There are many methods for obtaining absolute abundances from the relative abundances measured by MGS, but these methods have not previously been evaluated for their sensitivity to taxonomic bias (Williamson, Hughes, and Willis (2021) being a notable exception). Our theoretical results show that these methods differ in the extent to which taxonomic bias causes the FEs in species abundances to vary across samples. The effect of bias on absolute DA analysis can be mitigated by choosing methods with more stable FEs, using either of two general approaches.

#### 5.3.1 Use complementary MGS and total-abundance measurements

A popular approach to measuring the absolute abundance of individual species is to multiply the species’ proportions from MGS by a measurement of the abundance of the total community (Equation (11)). The resulting species abundances are affected by taxonomic bias in the MGS *and* the total-abundance measurement (Equation (14)). Section 3 shows that the effect of both forms of bias on DA results can be reduced by choosing MGS and total-abundance methods that have similar taxonomic bias.

Consider the debate over whether flow cytometry or 16S qPCR are better methods for measuring total abundance for the purposes of normalizing 16S amplicon sequencing data (Galazzo et al. (2020), Jian, Salonen, and Korpela (2021)). Flow cytometry directly counts cells, whereas 16S qPCR measures the concentration of the 16S gene following DNA extraction. Thus the 16S qPCR measurement is subject to bias from extraction, copy-number variation, primer binding, and amplification, just like the 16S sequencing measurement. Although this shared bias likely makes 16S-qPCR less accurate as a measure of total cell concentration, our theory suggests that it will lead to more accurate FDs—and thus more accurate DA results.

More generally, these observations suggest that for the purposes of performing an absolute DA analysis from amplicon sequencing measurements, the ideal total-abundance measurement is qPCR of the same marker gene, from the same DNA extraction. Similar reasoning suggests that for shotgun sequencing, the ideal total-abundance measurement is bulk DNA quantification: Shotgun sequencing and bulk DNA quantification are both subject to bias from extraction and variation in genome size. Optimal use of these pairings requires thoughtful choices during bioinformatics. For example, performing copy-number correction on amplicon read counts prior to multiplication by qPCR measurements would be counter-productive, as it decouples bias in the two measurements. Additionally, the MGS proportions should be computed prior to discarding any unassigned reads, since species that are missing from the given taxonomy database still contribute to the total concentration of marker-gene and/or bulk DNA. Future experiments should evaluate the extent to which the taxonomic bias of DNA-based total-abundance measurements is stable across samples and shared with the complementary MGS measurement.

#### 5.3.2 Normalize to a reference species

A second approach to obtaining absolute species abundances involves normalizing the MGS count for each species to that of a reference species with constant and/or known abundance (Equation (15)). Our theory predicts that taxonomic bias induces constant FEs in the abundances obtained by this approach, which will not affect DA results.

Previous studies have used species with a constant abundance as references: spike-ins, computationally-identified ‘housekeeping species’, and the host. Researchers may naturally be concerned about bias between the reference species and the native species being normalized against it (see, for example, Harrison et al. (2021)’s recommendations for spike-in experiments). Our results show that accurate FDs and DA results can be obtained so long as the relative efficiency of the focal species to the reference is consistent across samples. This condition is much weaker than requiring that bias is small (the relative efficiency is close to 1) and so greatly expands the applicability of spike-ins and host normalization in microbiome experiments.

Section 2.3 further proposed an additional class of reference species for normalizing MGS measurements: Species whose (absolute) abundance has been measured using a targeted method such as qPCR with species-specific primers. Direct, targeted measurement of a reference species removes the need for it to have a constant abundance, thus expanding the applicability of reference-species normalization to experiments where spike-ins or housekeeping species are not viable options. The targeted measurement need not itself be an unbiased measure of the reference species’ abundance; so long as it has constant FE, the abundances of native species obtained by normalization will also have constant FEs. To demonstrate the ability for reference calibration to reduce error in FD measurements, we treated the abundance of one species (*Lactobacillus crispatus*) in the Brooks et al. (2015) mock community data as known, and used it to calibrate the abundances of all species. Doing so improved the resulting measurements of FDs in the abundance of all species (Figure 4).

In some ecosystems, there may not be a single universally-present species that can serve as a reference in all samples. In such cases, sample coverage can be increased by performing targeted measurement of multiple species with complementary prevalence patterns, using appropriate statistical models to combine information across multiple measurements and impute the appropriate correction factor when all reference taxa are below the detection limit in the targeted or MGS measurement. Sample coverage can alternatively increased by targeting a larger taxonomic group (such as a genus or family); however, these higher-order taxa are less likely to have a consistent efficiency across samples (Section 2.2).

### 5.4 Bias sensitivity analysis

Even if control measurements are not available, it is possible to computationally assess the sensitivity of a given DA result to taxonomic bias. A *bias sensitivity analysis* can be conducted by analyzing an MGS dataset multiple times under a range of possible taxonomic biases. First, a large number of hypothetical biases are generated from a user-specified probability distribution. Next, the DA analysis is re-run while using each simulated bias vector to calibrate the MGS measurements. This approach is highly flexible, and can be used to investigate the bias sensitivity of any DA method. Alternatively, existing DA methods can be extended to directly include the unknown taxonomic bias in their underlying statistical model, thereby providing DA estimates that inherently account for the added uncertainty in microbiome compositions due to the presence of unknown bias. (Greenland et al. (2005) compares these two approaches in a general context; Nixon et al. (2022) applies the second approach to microbiome data in the absence of taxonomic bias.)

Interpreting a bias sensitivity analysis is complicated by the large percentage of zero counts in species-level microbiome profiles, as the results may strongly depend on how these zeros are handled in the calibration process. Therefore it can be advantageous to jointly test the sensitivity of assumptions about zero-generating processes along with taxonomic bias, making statistical DA models that include bias and one or more zero-generating processes especially valuable. The development of tools and workflows to facilitate bias sensitivity analysis may provide an efficient way to increase scientists’ ability to assess the reliability of microbiome results, both for differential abundance and microbiome analyses more generally.

### 5.5 Bias-aware meta-analysis

Meta-analysis of microbiome samples measured across multiple studies must contend with the fact that different studies typically use different protocols and hence have different taxonomic biases. These different biases can be explicitly accounted for in a *bias-aware meta-analysis* which has the potential to improve statistical power as well as interpretability of multi-study DA analyses. Parametric meta-analysis models include study-specific latent parameters representing ‘batch effects’—non-biological differences in the data from each study which can distort the observed biological patterns. By estimating these ‘nuisance parameters’ along with the biological parameters of interest (such as the difference in a species’ log abundance between conditions), the meta-analysis aims to reduce statistical bias in the biological parameters created by the non-biological differences among studies (Leek et al. (2010),Wang and LêCao (2020)). In a bias-aware meta-analysis, the meta-analysis model is configured so that (some of) the latent parameters correspond to study- and species-specific relative efficiencies. If taxonomic bias is consistent within but not between studies, then this approach may improve the ability for the meta-analysis to accurately identify DA patterns as compared to meta-analysis methods in which the ‘batch effects’ do not reflect the multiplicative and compositional nature of taxonomic bias. The ability to infer and adjust for the differential bias between studies can be increased by measuring one or more shared control samples alongside the experiment-specific samples.

## 6 Conclusion

It is commonly assumed that differential abundance (DA) analysis is robust to taxonomic bias, so long as all samples have been subjected to the same MGS workflow. In contrast, our results show that consistent taxonomic bias can distort the results of DA methods that are based on species proportions. This distortion occurs because the fold error in a species’ proportion depends on the sample mean efficiency, which can vary across samples and conditions in accordance with the overall community composition. This problem continues to apply for analyses of absolute abundance that are based on multiplying the proportions from MGS by the total abundance of the community. In contrast, DA methods based on species ratios are invariant to consistent bias, since the error in the ratio between species is independent of community composition. In Section 5, we describe how ratio-based methods can thus provide a more robust approach to analyzing relative and absolute abundances; in addition, calibration with community controls, bias-sensitivity analysis, and bias-aware meta-analysis provide additional ways to correct or mitigate the effect of bias.

Proportion-based DA methods encompass many of the most popular methods for analyzing relative and absolute abundances. Importantly, however, if the mean efficiency is approximately constant across samples, then bias has a negligible effect on multiplicative and rank-based DA results. If the mean efficiency varies but is unassociated with the condition of interest, then bias merely serves to increase noise and does not create systematic errors in DA results. It is only when the mean efficiency is associated with the condition being analyzed that large systematic errors can occur. Our case studies suggest that this problematic scenario does occur, but it may be the exception rather than the rule. Systematic investigation of how the mean efficiency affects DA results across a wide range of studies may increase our confidence in previous DA results and/or alert us to the conditions in which they are most suspect.

Important open questions include 1) determining the extent to which bias is consistent across samples for different taxonomic levels, MGS methods, and sample types; 2) assessing the validity of post-extraction measurements as measures of pre-extraction abundance; 3) understanding how the multiplicative error from bias interacts with the non-multiplicative error from contamination and taxonomic misassignment; and 4) understanding how different underlying community dynamics, and in particular the source of zero counts in MGS measurements, affect bias sensitivity. In addition, while we have showed some simple methods for incorporating bias and control measurements into DA analyses, more sophisticated statistical tools are needed to properly account for both taxonomic bias and random variation in the underlying MGS and supplemental (e.g. qPCR) measurements. Finally, more concrete experimental recommendations and user-friendly statistical workflows are needed for implementing the solutions we propose in various experimental contexts.

Our theoretical framework and example analyses provide a foundation for addressing these open questions, as well as developing experimental protocols and statistical tools that implement our proposed solutions. We look to a future in which microbiome researchers regularly choose an appropriate combination of experimental and data-analytic methods that are capable of answering their fundamental question while also accounting for taxonomic bias and other limitations inherent to MGS measurement. In doing so, we will collectively gain the confidence needed to codify the findings from MGS-based microbiome studies into true scientific knowledge.

## A Linear regression

## A.1 Simple linear regression

This section derives the theoretical results of Section 3 for the effect of taxonomic bias on regression-based DA analyses. As in Section 3, we restrict our attention to DA analyses that can be expressed in terms of the simple linear regression model in which the response is log absolute abundance or log proportion of individual species or the log ratio of a pair of species.

## A.1.1 Review of simple linear regression

We begin by reviewing definitions, notation, and results for the analysis of the simple linear regression model. Our presentation follows Wasserman (2004) Chapter 13 with additional interpretation of the estimated regression coefficients. Statistical notation is defined in Table 1.

**Table 1:**
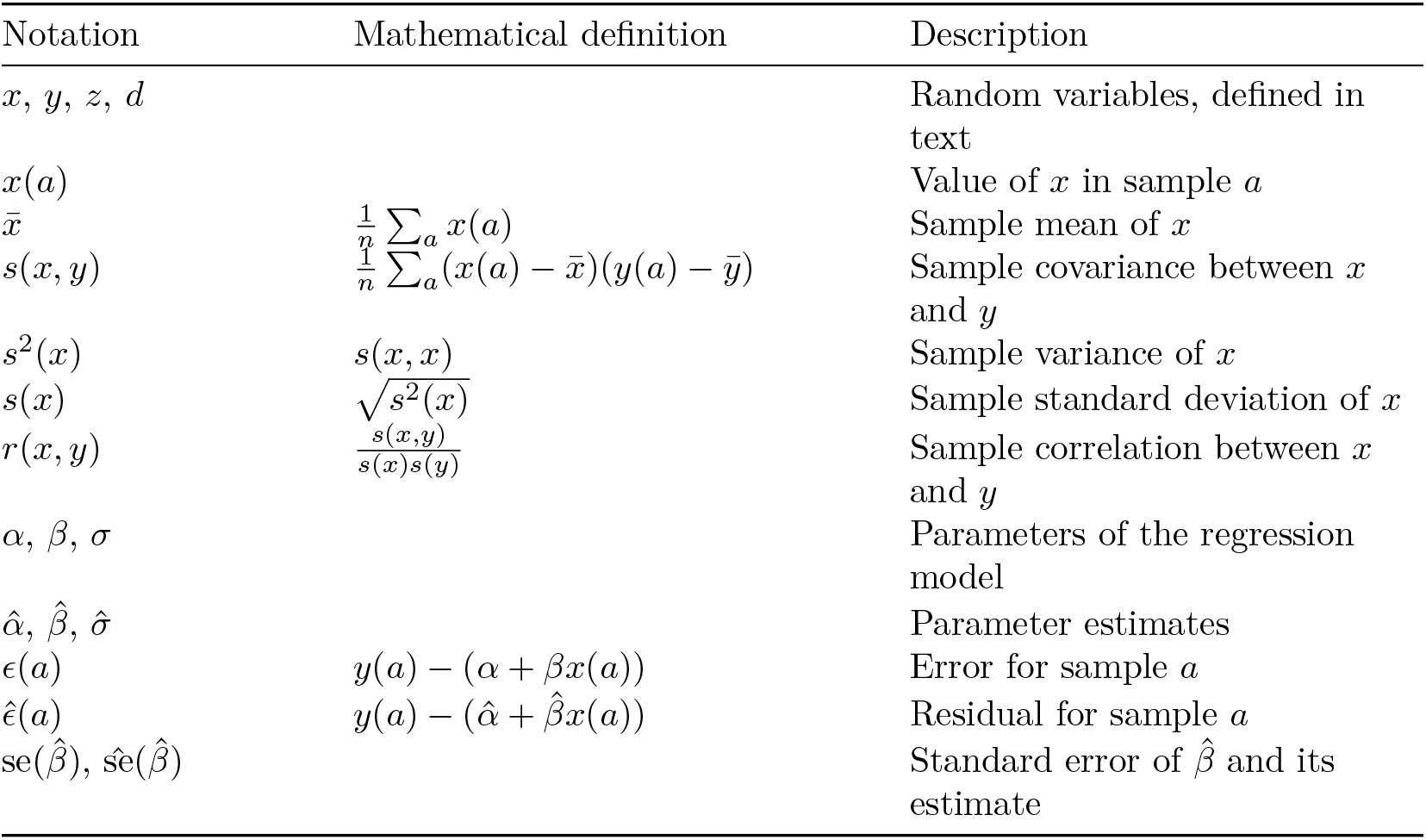
Statistical notation used in this section.

Consider a response variable *y* and a covariate *x*. Following Wasserman (2004), we define the *simple linear regression model* by

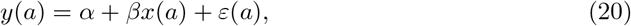

where *E*[*ε*(*a*) | *x*(*a*)] = 0 and *V* [*ε*(*a*) | *x*(*a*)] = *σ*^2^. That is, conditional on the value of *x* in a sample *a*, the response *y* is given by *α* + *βx* with a random error *ε* that has an expected value of 0 and a constant variance *σ*^2^.

Given data (*y, x*), we *fit* the model by finding estimates for the parameters *α*, *β*, and *σ*^2^ that best reflect the data under the assumption that the model is correct and according to our chosen criterion that defines ‘best’. Here we consider the least-squares estimates for the coefficients *α* and *β*, which are also the maximum likelihood estimates if we further assume that the errors *ε* are normally distributed. The least-squares estimates of *α* and *β* are the values that minimize the residual sum of squares, 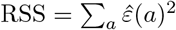. These estimates can be conveniently expressed in terms of the sample means and covariances or correlations of *x* and *y*,

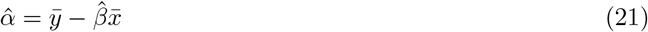

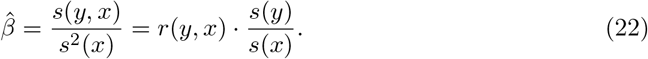

The estimate for the slope coefficient, 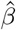, equals the ratio of the sample covariance of *y* with *x* relative to the variance in *x* or, equivalently, the correlation of *y* with *x* multiplied by the ratio of their standard deviations. The estimate for the intercept, 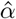, is the value such that the sample means follow the regression relationship, 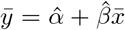. The standard unbiased and maximum likelihood estimates for the variance *σ*^2^ are both approximately given by the sample variance of the residuals,

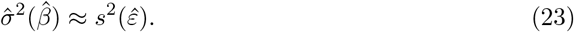

We are most interested in the estimated slope coefficient, 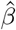. Besides the point estimate indicating our best guess at the value, we also wish to understand the uncertainty or precision of the estimate. This uncertainty is quantified by the standard error, 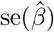, estimated by

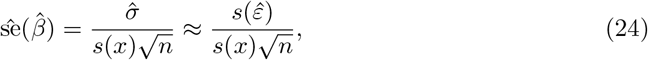

Approximate 95% confidence intervals for *β* are given by 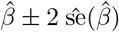.

## A.1.2 Measurement error in the response

Now consider a researcher who would like to regress a biological quantity *y* (the response) on a second quantity *x* (the covariate). But instead of *y*, they only know a proxy *z*, which is related to *y* via the difference *d* = *z* − *y*. For instance, *y* might be the log abundance of a species, *z* is the abundance that is measured via MGS, and *x* is a numerical quantity like pH or a boolean variable indicting case versus control. Lacking a direct measurement of *y*, the researcher instead regresses *z* on *x* and interprets the fitted model as informing them about the relationship between *y* and *x*. We wish to understand the accuracy of their conclusions.

To understand how the measurement error in *y* impacts the researcher’s regression analysis, consider the three linear regression equations for *y*, *z*, and *d*,

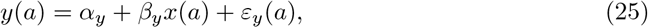

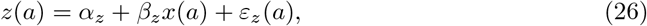

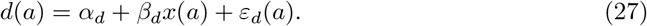

The researcher would like to fit the model for *y* (Equation (25)), but instead can only fit the model for *z* (Equation (26)). The two are related via the Equation (26) for *d*. It is helpful to imagine that *y* and *d* are known and so we can fit all three models (25), (26), and (27). From Equation (21) and the linearity of sample means and covariances it follows that

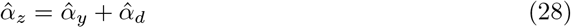

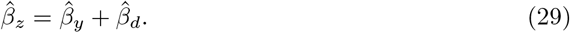

That is, the estimated coefficients for *y*, *z*, and *d* mirror the relationship between the variables themselves, *z* = *y* + *d*. Consequently, we see that the measurement error *d* creates absolute errors in the regression coefficients 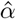 and 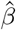 equal to

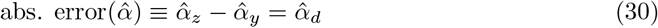

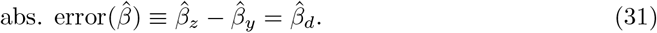

Expressed in terms of sample means and covariances, these errors are

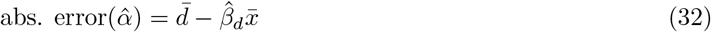

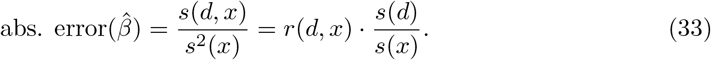

We are mainly interested in the estimated slope coefficient, 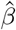. We see that the absolute error is large when the covariance of *d* with *x* is large. This error is large in a practical sense when 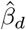 is large (in magnitude) relative to 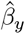 (Equation (21)), which occurs when *s*(*d, x*) is large relative to *s*(*y, x*). A sign error occurs when *s*(*d, x*) is larger in magnitude than *s*(*y, x*) and of opposite sign.

We can also consider how the measurement error affects the residual variation and the standard error of the slope estimate. The residuals of *y*, *z*, and *d* are related through 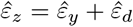. It follows that the sample variance of the *z* residuals equal

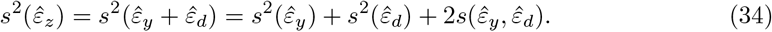

The variance of the *z* residuals is increased above that of the *y* residuals when the *d* residuals are either uncorrelated or positively correlated with the *y* residuals, but may be decreased when the *y* and *d* residuals are negatively correlated. An increased residual variance will lead to a larger estimate for *σ* as well as larger standard errors in the slope coefficient *β*.

## A.1.3 Specific application to taxonomic bias

These general results can be used to understand how taxonomic bias affects DA analyses that can be expressed as linear regression of log (relative or absolute) abundance.

First, we illustrate for the particular example of absolute DA analysis, where species absolute abundance is measured with the total-abundance normalization method and we assume that the total abundances have been accurately measured. Consider the regression of log abundance of species *i* on a covariate *x*. In this case, *y*(*a*) = log abun_*i*_(*a*) is the actual log abundance of species *i* and 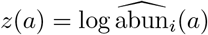 is the log abundance we’ve measured. From Equation (14), the measurement error *d*(*a*) due to taxonomic bias in the MGS measurement is

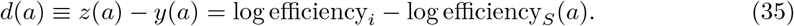

The absolute error in 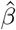 equals the scaled covariance of *d* with *x* (Equation (32)). In the above expression for *d*(*a*), only the second term, −log efficiency_*S*_(*a*), varies across samples and thus affects the covariance, leading to

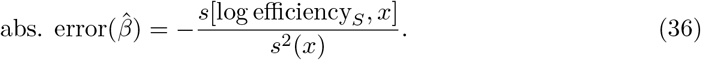

Thus the absolute error in 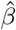 is the negative of the scaled covariance of the log mean efficiency.

Although the absolute error is the same for each species, its practical significance varies. Recall from Equation (21) that the correct value for 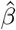 (i.e., the estimate without measurement error) is

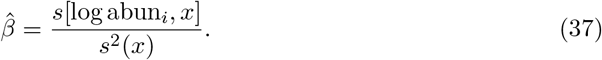

For species whose log abundance covaries with *x* much more than the log mean efficiency, the error will be relatively small. This situation can occur either because the mean efficiency varies relatively little across samples or because its variation is relatively uncorrelated with *x*.

For species whose log abundance covaries with *x* similar or less than the log mean efficiency, the error will be significant. A sign error occurs when the species covariance is of *equal sign and smaller in magnitude* than the covariance of the log mean efficiency.

We can also consider how the standard errors in the slope estimate are affected by bias. In our case, the *d* residuals equal the negative residuals of the log mean efficiency. It is plausible that for most species, their residual variation will have a small covariance with log mean efficiency and the net effect of variation in the mean efficiency will be to increase the estimated standard errors, as occurs with most species in Figure 2. However, high-efficiency species that vary substantially in proportion across samples may be strongly positively correlated with log mean efficiency such that the estimated standard errors decrease, as we see with Species 9 in Figure 2.

Now, suppose we were instead performing a DA analysis of a response whose fold error is constant across samples, such as the log ratio of two species. For the log ratio of species *i* and *j*, the error is *d*(*a*) = log efficiency_*i*_/efficiency_*j*_ and is constant across samples. Thus the covariance of *d* with *x* is 0 and taxonomic bias causes no error the slope estimate 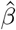, only the intercept estimate 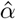.

## A.2 Gamma-Poisson regression

This section describes how gamma-Poisson regression can, with appropriate choice of offsets, be used to estimate log fold changes (LFCs) in proportions from MGS count data either with or without bias correction.

## A.2.1 Background

The gamma-Poisson distribution, also known as the negative binomial distribution, is a distribution that is commonly used to model sequencing count data (Holmes and Huber (2018)). Its key advantage over the simple linear regression model is that it directly models the random sampling of reads that occurs during a sequencing experiment and, in this way, naturally accounts for the imprecision associated with an observation of zero or a small positive count for a given species and sample. The gamma-Poisson distribution has two parameters, which jointly determine the mean and the standard deviation. We use the parameterization used by the rstanarm R package (Goodrich et al. (2020)), which we use for fitting gamma-Poisson GLMs in our case-study analyses. Suppose that *y*^(*a*)^ has a gamma-Poisson distribution conditional on a covariate *x*. Define two parameters *μ* and *ϕ* such that

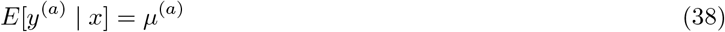

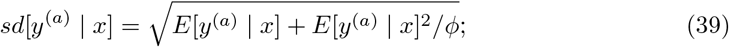

thus *μ* equals the mean and *ϕ*, known as the ‘reciprocal dispersion’, increases the standard deviation as it approaches 0 and causes the standard deviation to approach 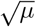 (that of a Poisson distribution) as it approaches infinity.

For use in gamma-Poisson GLM regression of MGS data, we partition the mean into two factors, *μ*^(*a*)^ = *u*^(*a*)^*θ*^(*a*)^ (Gelman, Hill, and Vehtari (2020)). Here, 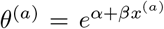 is the quantity of primary interest, whose average LFC equals *β*. The factor *u*^(*a*)^ is a sample-specific *exposure* parameter that determines the overall scale of counts. We can equivalently write 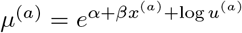; the logarithm of the exposure, log *u*^(*a*)^, is known as the *offset* of the GLM. Performing gamma-Poisson regression with standard statistical software requires that the offsets are provided as known values.

## A.2.2 Inferring LFCs in proportions with and without bias correction

Our goal is to use gamma-Poisson regression to estimate the LFC in a species proportion from the observed read counts. We therefore equate the counts 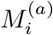 with *y*^(*a*)^ and the proportion 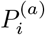 with *θ*^(*a*)^; it remains to determine the offsets that are consistent with our MGS model. To do so, we relax the deterministic assumption of our original model and instead suppose that the right-hand side of Equation (1) equals the *expected* read count of species *i* in sample *a*,

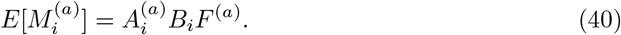

The total expected count is

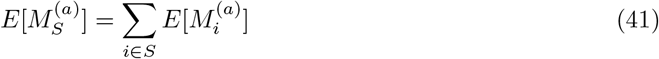

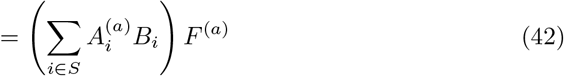

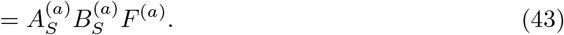

Under the gamma-Poisson model for 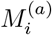, we have that 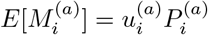. Equating this expression for 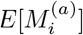 with Equation (40) lets us solve for the exposure,

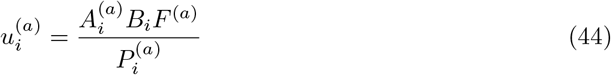

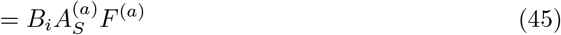

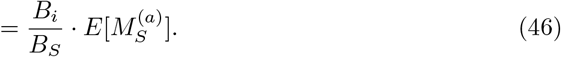

The second line follows from 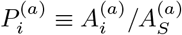, and the third line from Equation (41).

The final expression in Equation (44) can be used to compute offsets for the regression. The three terms *B_i_*, 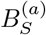, and 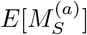 are each unknown, so we instead substitute estimates for these terms to obtain an estimated exposure 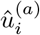; we then set the offset in the linear model to 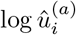. We can estimate 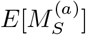 by the observed total count, 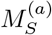. In the absence of bias correction, we set 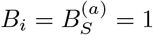; our estimate of the exposure is then just

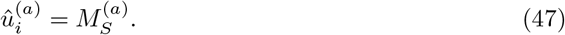

We can apply bias correction using estimates of the efficiencies, 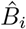, derived from community control measurements. From these estimates and the observed counts, we can estimate the mean efficiency in the sample by

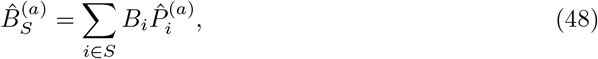

where 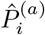 are the calibrated proportions obtained from a simple plug-in procedure,

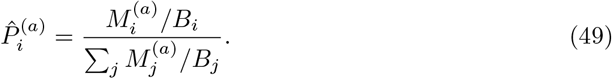

Equivalently, we can calculate 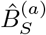 directly from the observed counts,

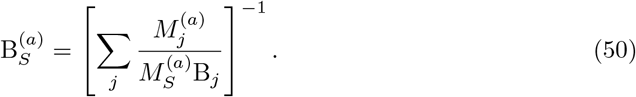

We then estimate the exposure by

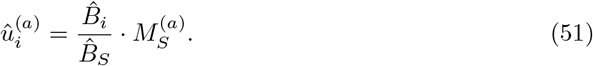

## B Supplemental figures

**Figure 5:**
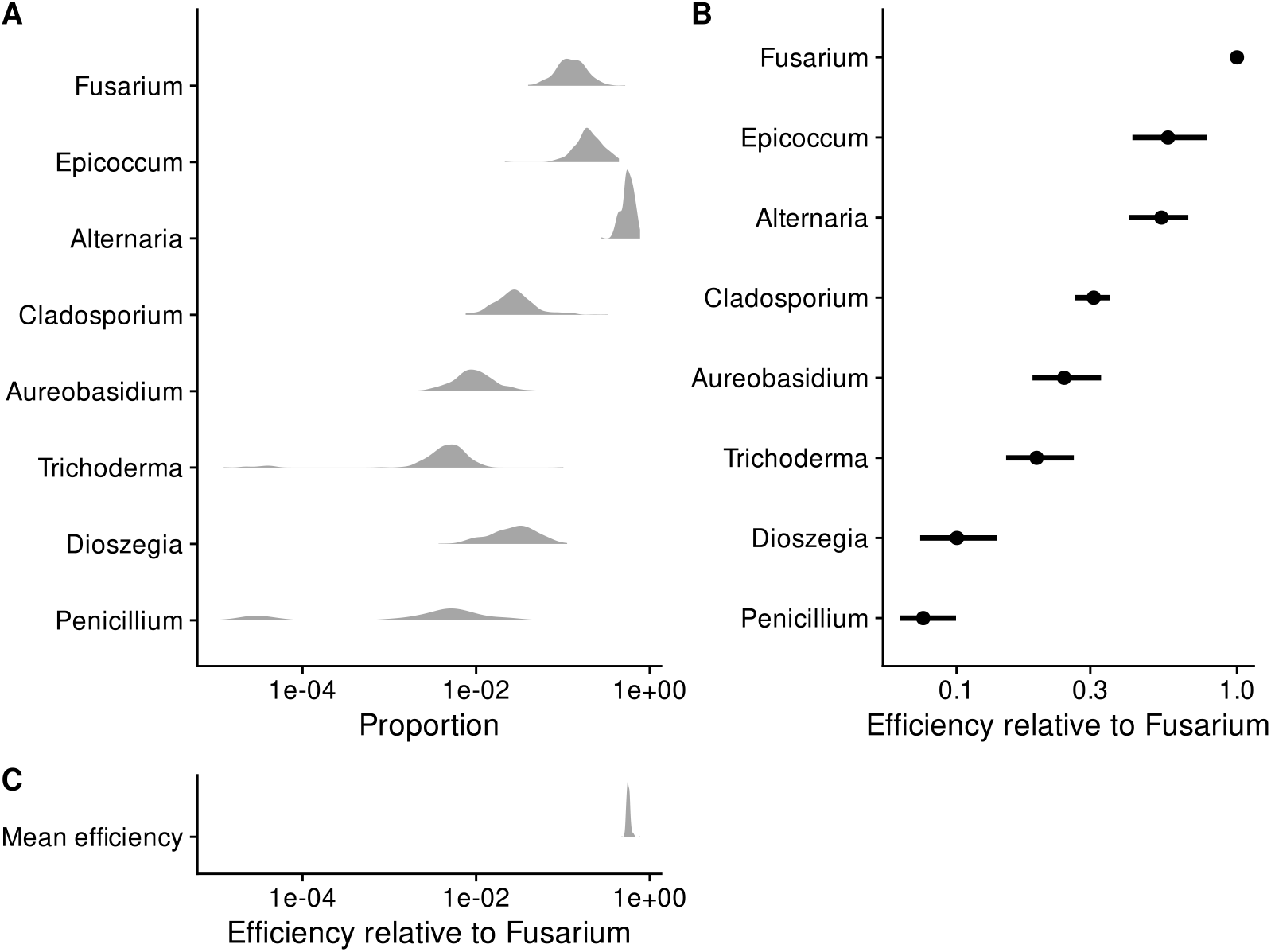
In the pre-infection samples from Leopold and Busby (2020), multiplicative variation in taxa proportions is much larger than that in the mean efficiency. Panel A shows the distribution of the proportions of each commensal isolate (denoted by its genus) across all samples collected prior to pathogen inoculation; Panel C shows the distribution of the (estimated) sample mean efficiency across these same samples on the same scale; and Panel B shows the efficiency of each taxon estimated from DNA mock communities as point estimates and 90% bootstrap percentile confidence intervals. Efficiencies are shown relative to the most efficiently measured taxon (*Fusarium*).

**Figure 6:**
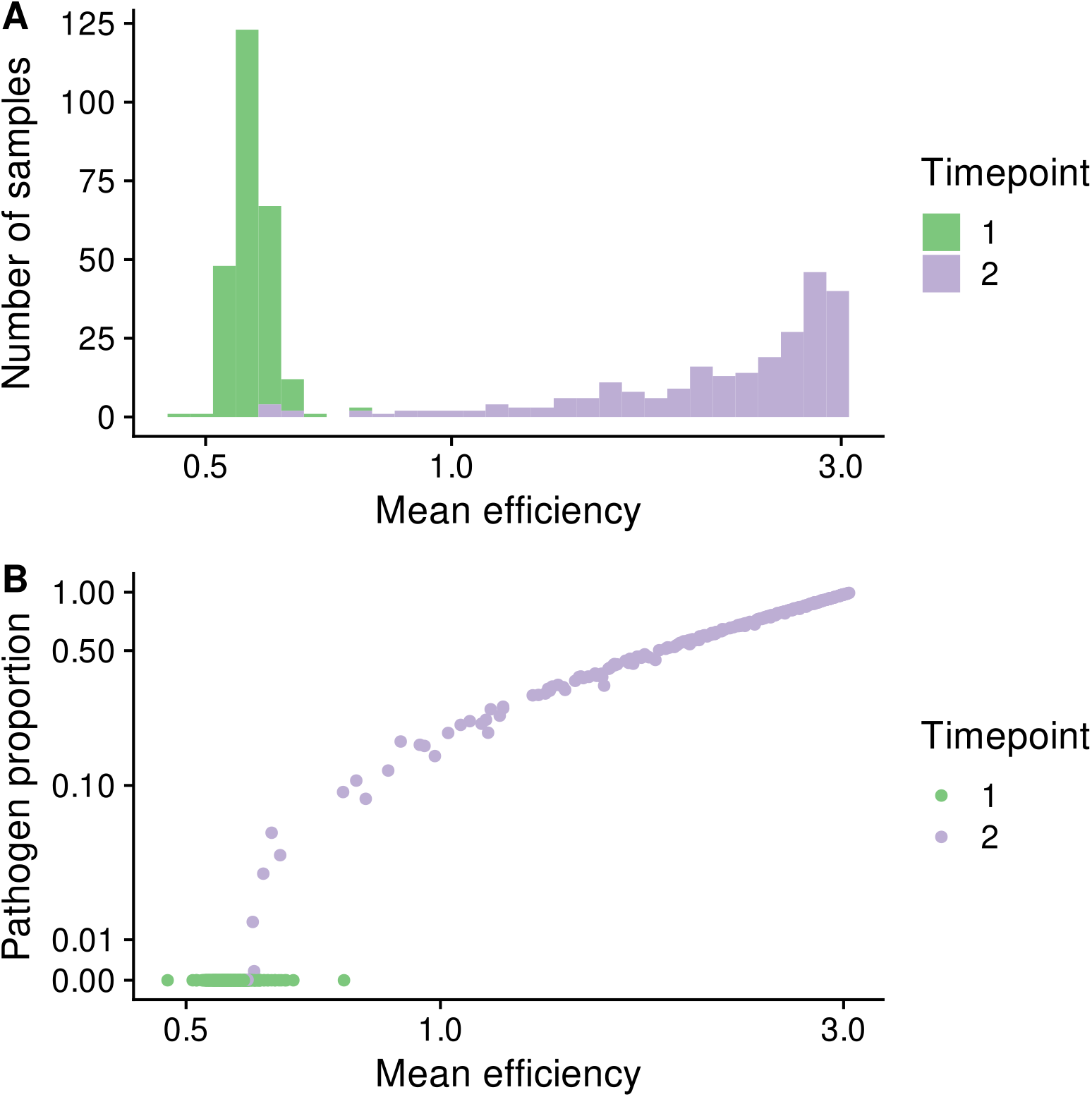
The mean efficiency tends to increase after infection due to the high proportion of the pathogen.

**Figure 7:**
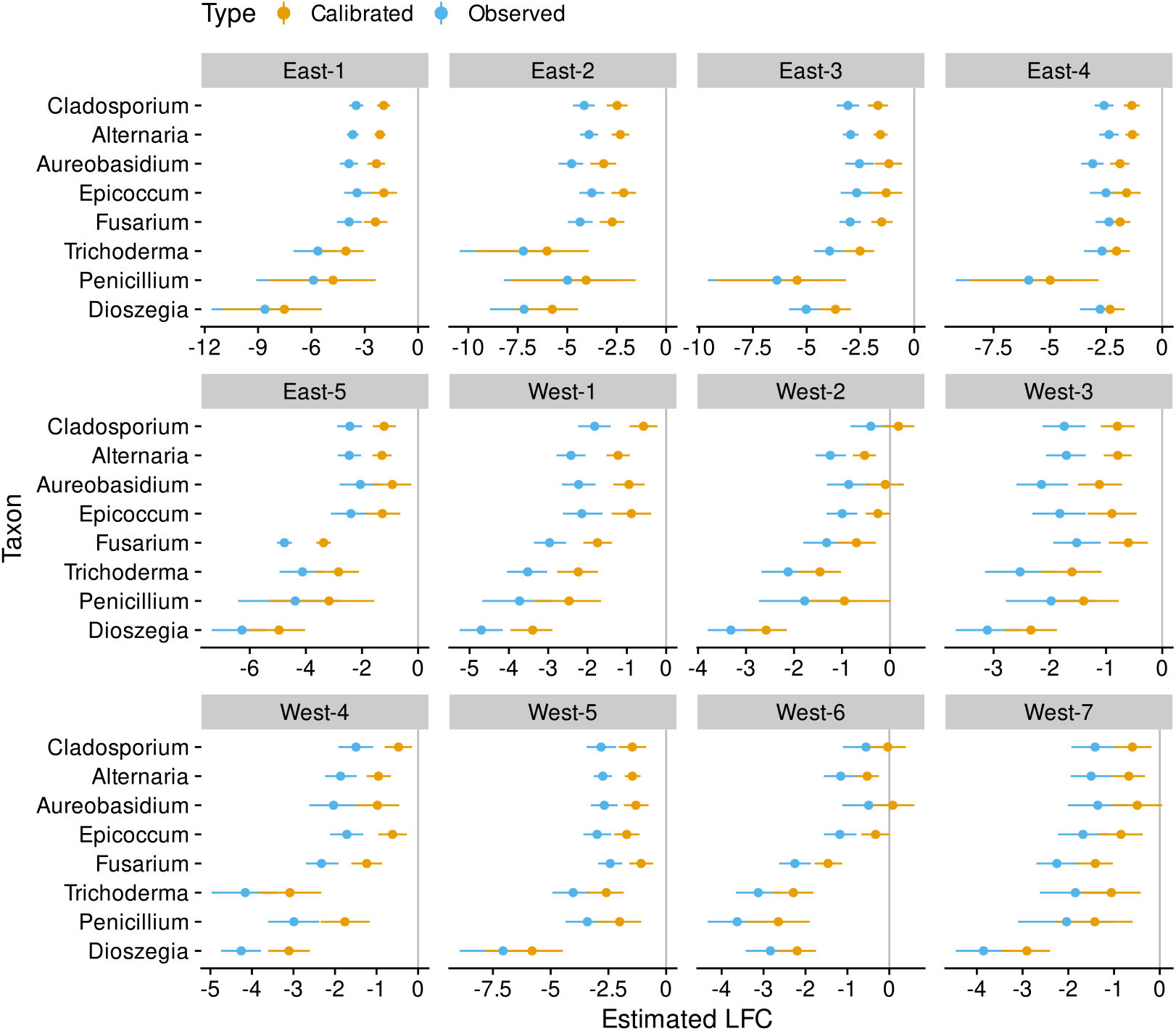
Bias correction increases the estimated log fold changes (LFCs) in commensal species proportions in response to pathogen infection. Shown are the estimated LFCs with 95% Bayesian credible intervals (CIs) derived from a Bayesian gamma Poisson regression. Negative values indicate that the species’ proportion decreased on average in response to infection, which is the default expectation given growth of the pathogen and the sum-to-one constraint of proportions. Bias leads to artificially low estimates, as the increase in the pathogen proportion increases the mean efficiency. In Western genotypes, several species whose proportion appears to decrease are instead found to remain stable or even to increase when bias is corrected.

**Figure 8:**
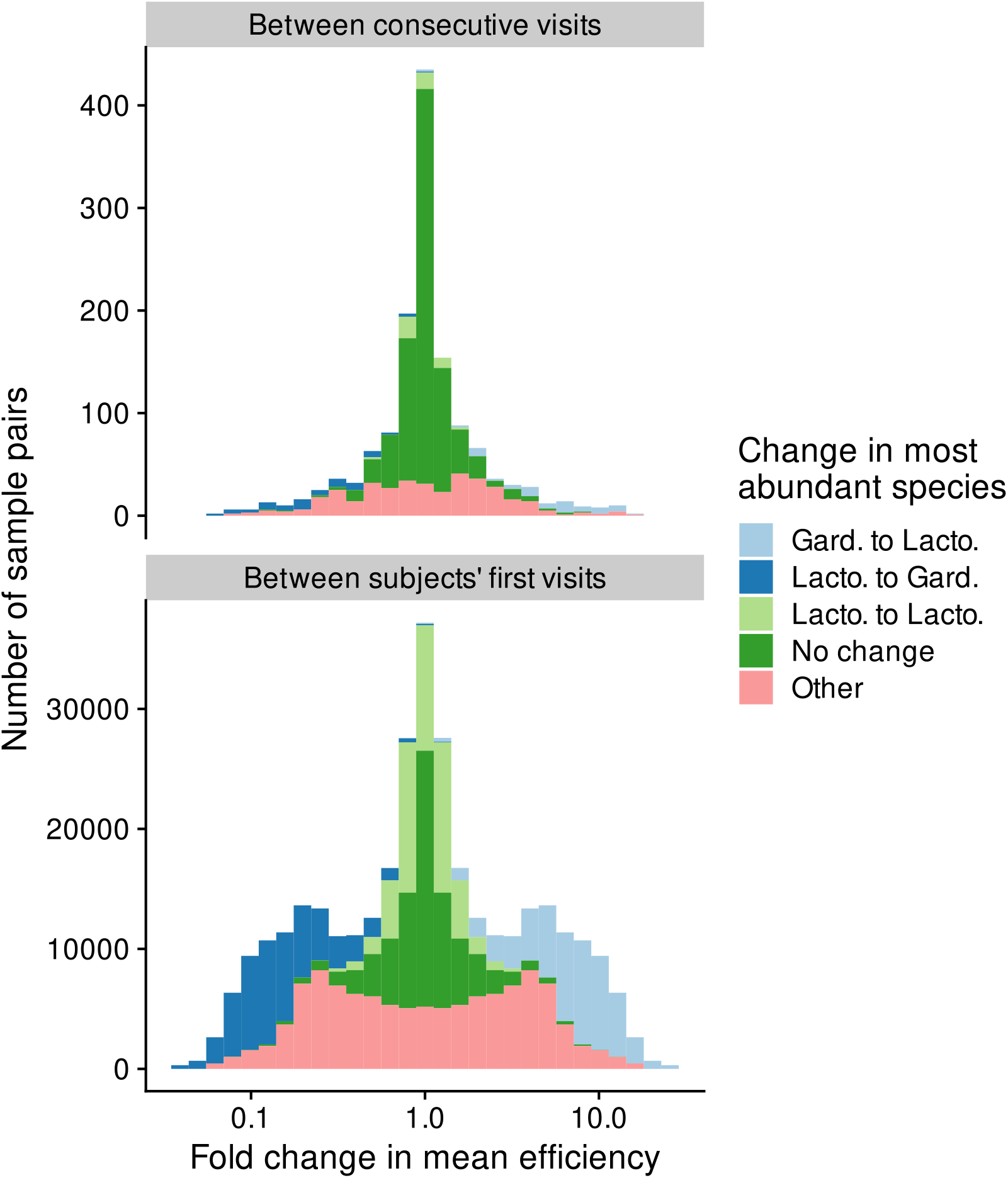
Fold changes in the mean efficiency within and between women in the MOMS-PI study.

**Figure 9:**
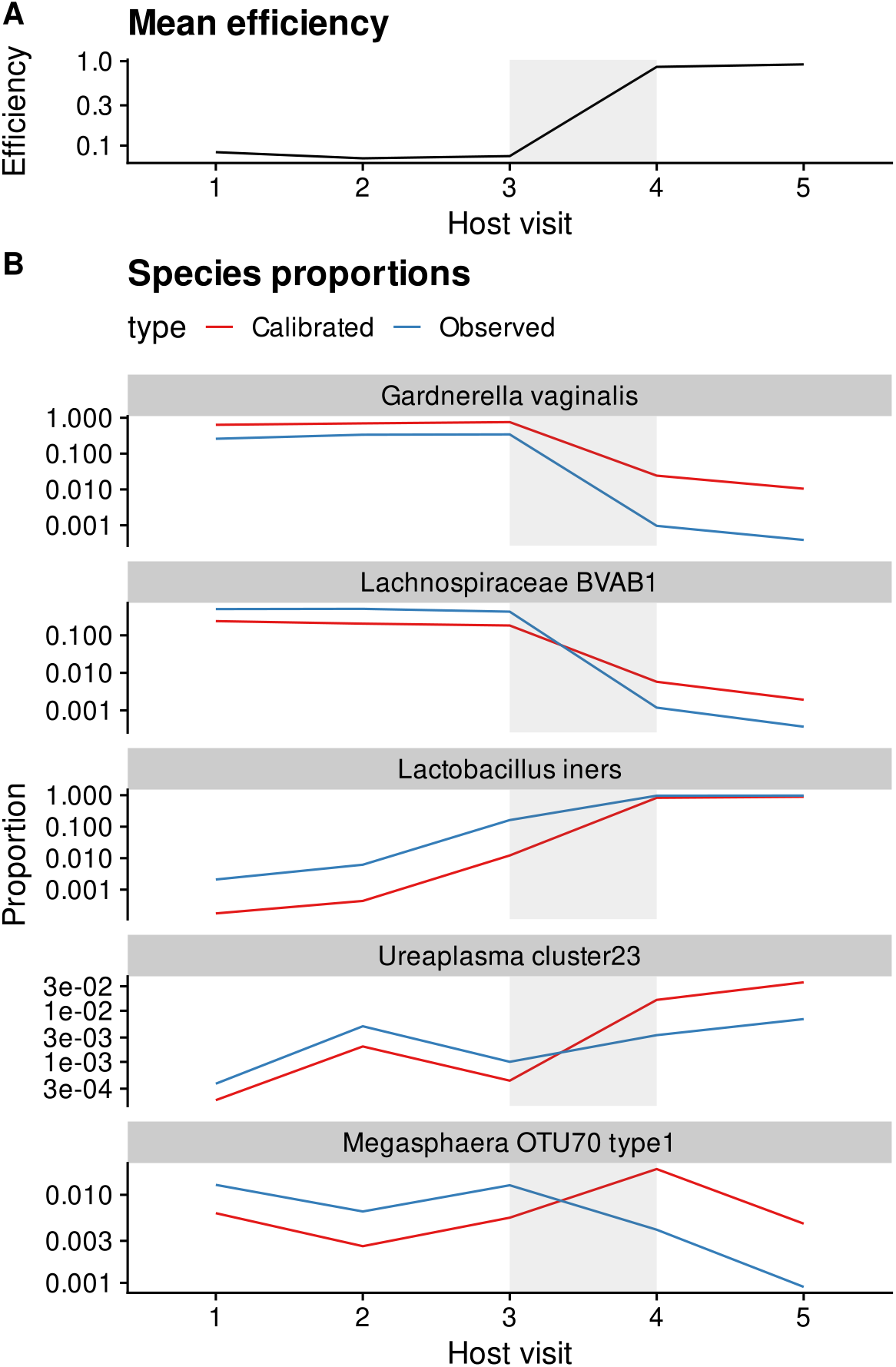
In vaginal microbiome measurements, shifts between *Lactobacillus* and *Gardnerella* dominance can drive spurious fold changes in other, lower-abundance species. The figure shows species proportions and mean efficiency trajectories over consecutive clinical visits for a subject in the MOMS-PI study whose microbiome samples showed substantial variation in mean efficiency. The subject’s samples are dominated by *Gardnerella vaginalis* and *Lachnospiraceae BVAB1* during the first three visits before transitioning to being dominated by *Lactobacillus iners* between visits 3 and 4. This transition drives a sharp increase in the mean efficiency, which significantly distorts the fold changes in the observed (uncalibrated) microbiome measurements for species with less dramatic fold changes. Two exemplar species are shown to illustrate the magnitude (*Ureaplasma cluster 23*) and direction (*Megasphaera OTU70 type1*) errors that can arise in this situation.

**Figure 10:**
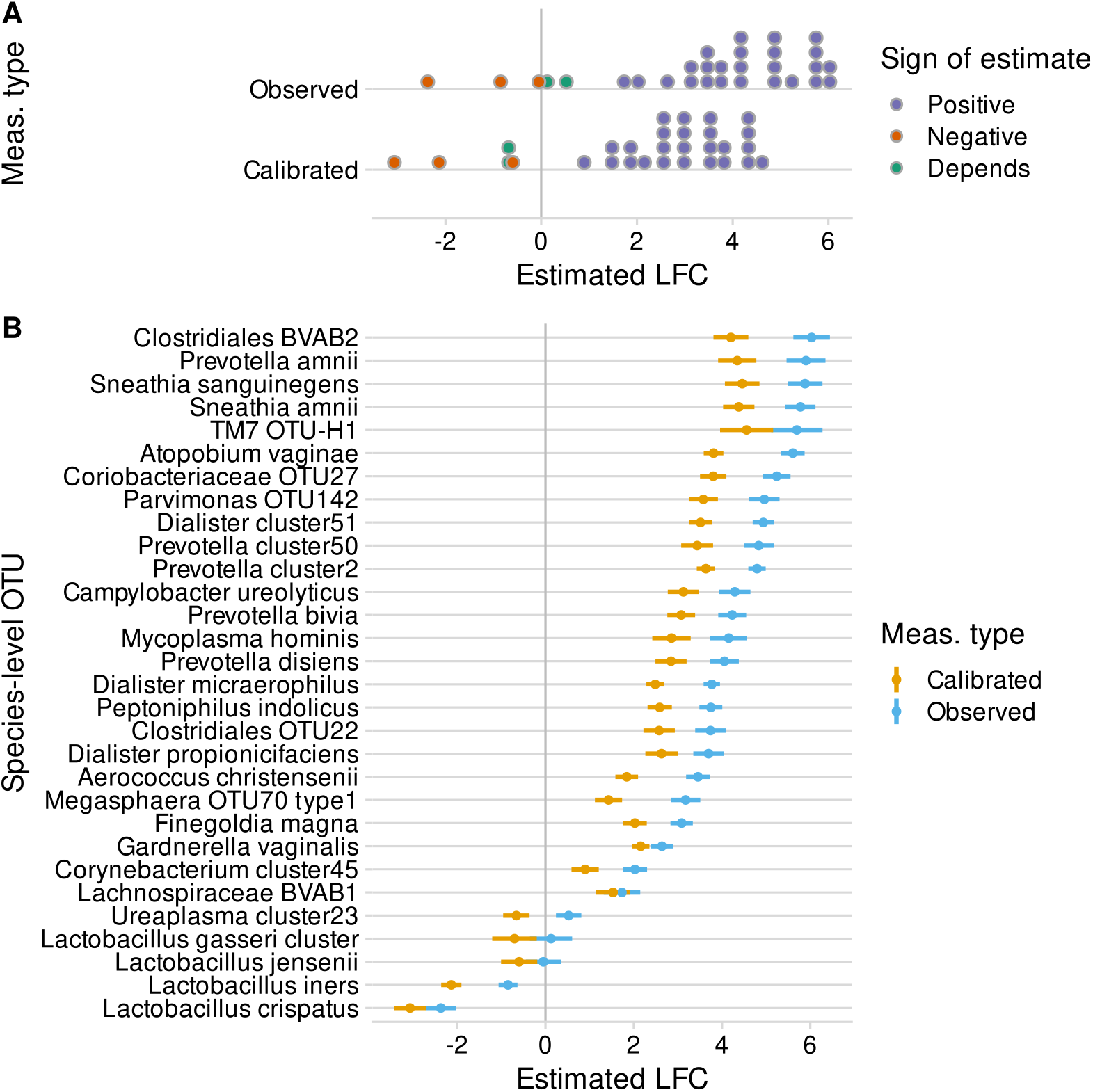
Bias inflates estimated log fold changes (LFCs) in species proportions with diversity in vaginal microbiome samples from the MOMS-PI study. Samples were split into low, medium, and high diversity groups based on Shannon diversity in observed (uncalibrated) microbiome profiles. The LFC in proportion from low- to high-diversity samples was estimated for 30 common species using gamma-Poisson regression with and without bias correction. Panel A shows the distribution of point estimates; Panel B shows the point estimates and 95% Bayesian credible intervals for each species.

**Figure 11:**
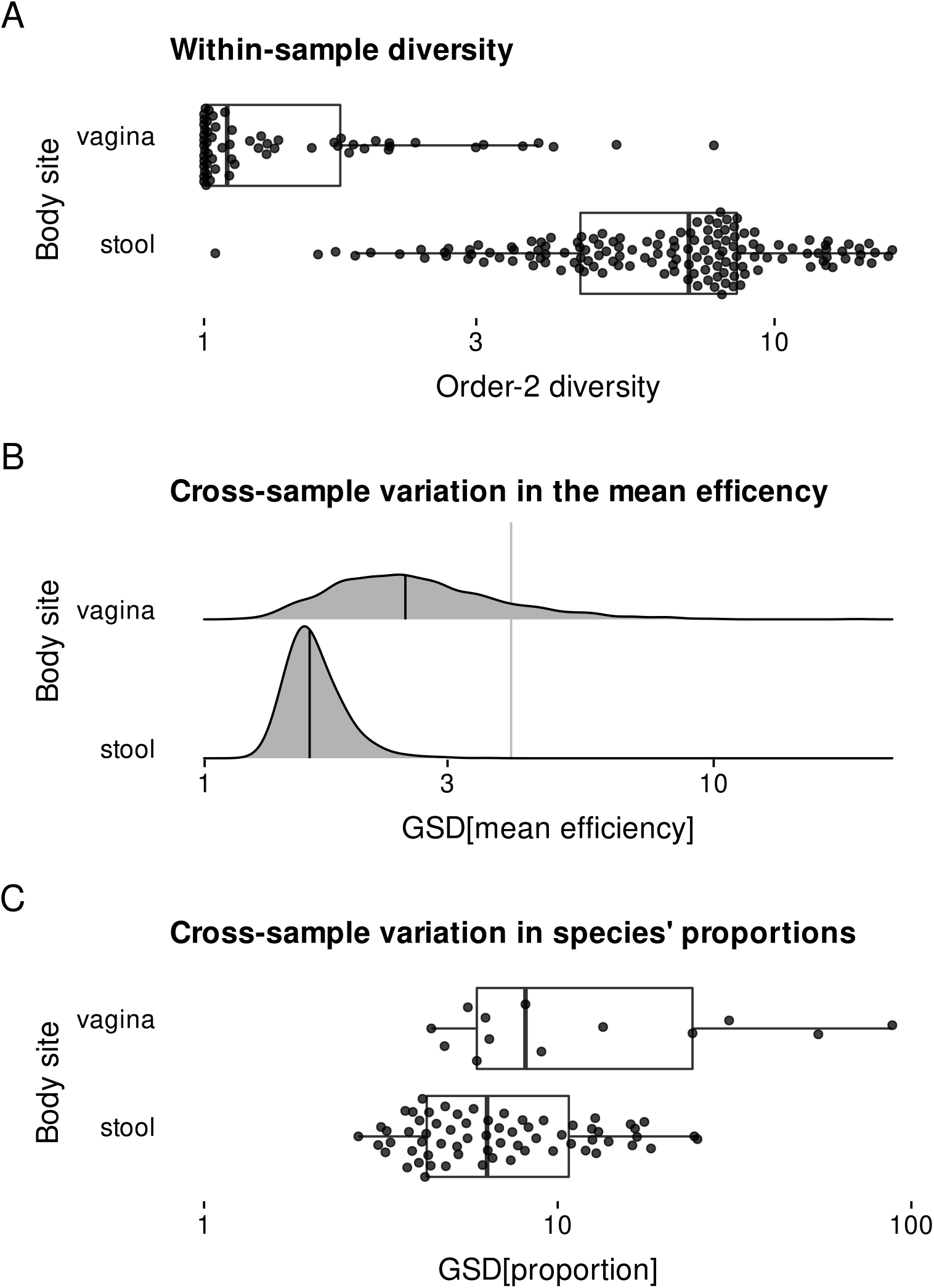
Compared to vaginal samples, stool samples have higher diversity, lower variation in mean efficiency, and lower variation in species proportions. This figure summarizes results from our case study of shotgun-sequenced stool and vaginal samples in the Human Microbiome Project. Panel A shows the distribution of order-2 alpha diversity (equal to the Inverse Simpson Index) across samples. Panel B shows the variation in the mean efficiency, as quantified by the GSD, across samples of a given type for each of 1000 simulated bias vectors. Panel C shows the variation in the proportion of each species commonly found in a given sample type. For Panel C, variation is measured by the GSD when a pseudo-value of 10^−4^ is added to the raw proportions. Different choices of the pseudo-value substantially change the measured GSDs, but not the difference (ratio) between gut and vaginal species.

## Acknowledgments

We thank Jen Nguyen and members of the Callahan Lab for valuable discussions.

